# Tyrosyl-DNA phosphodiesterase 1 (TDP1) and SPRTN protease repair histone 3 and topoisomerase 1 DNA-protein crosslinks *in vivo*

**DOI:** 10.1101/2023.03.01.530659

**Authors:** Ivan Anticevic, Cecile Otten, Luka Vinkovic, Luka Jukic, Marta Popovic

**Author notes:** Correspondence: Marta Popovic Corresponding Author.

## Abstract

DNA-protein crosslinks (DPCs) are frequent and damaging DNA lesions that affect all DNA transactions, which in turn can lead to the formation of DSBs, genomic instability and cell death. At the organismal level, impaired DPC repair (DPCR) is associated with cancer, aging, and neurodegeneration. Despite the severe consequences of DPCs, little is known about the processes underlying repair pathways at the organism level. SPRTN is a protease that removes most cellular DPCs during replication, whereas tyrosyl-DNA phosphodiesterase 1 repairs one of the most abundant enzymatic DPCs, topoisomerase 1-DPC (TOP1-DPC). How these two enzymes repair DPCs at the organism level is currently unknown. We perform phylogenetic, syntenic, structural and expression analysis to compare TDP1 orthologs between human, mouse and zebrafish. Using the zebrafish animal model and human cells, we demonstrate that TDP1 and SPRTN repair endogenous, camptothecin- and formaldehyde-induced DPCs, including histone H3- and TOP1-DPCs. We show that resolution of H3-DNA crosslinks depends on upstream proteolysis by SPRTN and subsequent peptide removal by TDP1 in RPE1 cells and zebrafish embryos, whereas SPRTN and TDP1 function in different pathways in the repair of endogenous TOP1-DPCs and total DPCs. Furthermore, our results suggest that TDP2 could potentially compensate for the impairment of TDP1 function *in vivo* and in human cells. Understanding the role of TDP1 in DPC repair at the cellular and organismal levels could provide an impetus for the development of new drugs and combination therapies with TOP1-DPC inducing drugs.

## Introduction

DNA-protein crosslinks (DPCs) are very frequent DNA lesions that are triggered by byproducts of normal cellular processes such as aldehydes, reactive oxygen species, and helical DNA modifications including apurinic/apyrimidinic (AP) sites (Vaz et al., 2017). They occur endogenously at a high frequency, with current estimates of 6000 lesions per cell per day (Ruggiano & Ramadan, 2021b). DPCs can also arise from the exposure to exogenous sources such as UV light, ionizing radiation, and chemotherapeutic agents (Fielden et al., 2018). These bulky lesions vary considerably depending on the type of DNA-binding protein, binding chemistry, and DNA topology near the lesion (single- and double-strand breaks or intact DNA). Any protein in the vicinity of DNA can be crosslinked upon exposure to the above triggers. Histones, high mobility group (HMG) proteins, and DNA-processing enzymes such as topoisomerases, DNA (cytosine-5)-methyltransferase 1 (DNMT1), poly [ADP-ribose] polymerase 1 (PARP1), and Ku70/80 have recently been identified as most common endogenous cellular DPCs (Kiianitsa & Maizels, 2020). Considering that DPCs affect all DNA transactions including replication, transcription, chromatin remodeling, and DNA repair, the consequences of impaired repair are severe and include single- and double-strand breaks, genomic instability, and apoptosis, which in turn can lead to cancer, accelerated aging and neurodegeneration. In contrast to other DNA repair pathways that have been extensively studied for decades, DPC repair is a relatively new field that was only recognized as a DNA Damage Repair (DDR) pathway in its own right after 2014, when the first protease that initiates DPC removal from the DNA backbone was found in yeast (Wss1) (Stingele et al., 2014).

DPCs can be repaired by proteolysis followed by removal of the peptide residues from the DNA backbone, or by nucleases that excise the part of the DNA to which the crosslinked protein is bound. The majority of cellular DPCs are thought to be removed by the SPRTN protease in dividing cells (Duxin et al., 2014; Maskey et al., 2017; Stingele et al., 2016; Vaz et al., 2016), while the exact contribution of other proteases, including ACRC/GCNA, FAM111A, DDI1, DDI2, and proteasome, remains to be determined (Ruggiano & Ramadan, 2021a). After proteolysis, the remaining crosslinked peptide could be repaired by the nucleotide excision repair pathway (Chesner & Campbell, 2018) or, in the case of crosslinked toposomerases 1 and 2 (TOP1- and 2-DPCs), by tyrosil-DNA phosphodiesterases 1 and 2 (TDP1 and 2) (Pommier et al., 2022).

Topoisomerases 1 and 2 create a transient DNA-protein intermediate to reduce DNA torsional stress. Camptothecin and its derivatives are used in cancer therapy to stabilize TOP1 intermediates and prevent replication of cancer cells (Fengzhi Li, 2017). TDP1 is a critical enzyme for TOP1-DPC repair and is a promising target for cancer treatment (Laev et al., 2016, Sun et al., 2020). The phosphotyrosyl bond between TOP1 and DNA is hydrolyzed by TDP1, releasing the TOP1 residue from the DNA backbone (Pommier et al., 2014). This bond is normally shielded and becomes accessible to TDP1 after partial proteolysis of TOP1. However, the dominant protease acting upstream of TDP1 has not yet been identified. While SPRTN has previously been considered the major protease for TOP1-DPC repair (Larsen et al., 2019; Reinking et al., 2020; Vaz et al., 2016), other proteases such as FAM111A, DDI1, and DDI2, as well as the proteasome, have also been associated with TOP1-DPC proteolysis (Kojima et al., 2020; C. P. Lin et al., 2008; Serbyn et al., 2020). Alternatively, another mechanism involving the 3’-flap endonucleases (Mus81/Eme1, Mre11/Rad50, and XPF/ERCC1) can facilitate TOP1-DPC removal without the need for proteolysis (Deng et al., 2005; Liu et al., 2002; Sun et al., 2020; Vance & Wilson, 2001, 2002). However, the nucleolytic pathway may not be as effective in cancer cells which often have mutated or inactivated DNA checkpoints required for endonuclease activation (Dexheimer et al., 2008), potentially making them more dependent on repair mediated by proteases and TDP1. Mutations in TDP1 and SPRTN lead to the neurological disorder spinocerebellar ataxia with axonal neuropathy-1 (SCAN1) (Takashima et al., 2002) and Ruijs–Aalfs syndrome (RJALS) characterized by premature aging and early-onset liver cancer (Lessel et al., 2014), respectively.

In addition to tyrosyl-3′-phosphodiester crosslink, TDP1 can resolve a variety of substrates at the 3’ end of DNA (Interthal et al., 2005; Raymond et al., 2004). Recently, it was shown that TDP1 can remove histones H2B and H4 attached to the AP sites in the *in vitro* system (Wei et al., 2022) without the need for prior proteolysis. However, TDP1 is unable to remove larger DPCs including Parp1 and TOP1 *in vitro* (Heidrun & James, 2011; Prasad et al., 2019; Wei et al., 2022). It is currently unknown how these discoveries translate to the *in vivo* system. Understanding the role of TDP1 in DPC repair at the cellular and organismal levels could provide an impetus for the development of new drugs and combination therapies with TOP1-DPC inducing drugs. Therefore, we have set out to investigate whether TDP1 can repair histone and TOP1-DPCs in cells and animal model and whether upstream proteolysis by SPRTN is required. Previously, it was thought that TDP1 specifically repairs TOP1-DPC remnants in cells, whereas data at the organism level were lacking. Our model organism, zebrafish (*Danio rerio*), is a well-established vertebrate systeml used to study cancer, neurodegenerative and cardiovascular diseases (Bradford et al., 2022; Choi et al., 2021). In zebrafish, DNA damage genes and pathways are 99% conserved compared to humans. Unlike mice, zebrafish are externally fertilized, which significantly facilitates gene editing, embryo manipulation, and DPC analysis. In addition, the very high fecundity allows for better statistical analysis of DPC formation and repair in both, embryos and adults (Choi et al., 2021; C. Y. Lin et al., 2016).

In this study, we investigate the function of TDP1 and SPRTN in the repair of TOP1- and histone H3-DPCs, as well as in the repair of total cellular DPCs in human cells and in zebrafish model. To this end, we compare human, mouse and zebrafish TDP1 orthologs using phylogenetic, syntenic, structural and expression analysis and show that zebrafish is a suitable vertebrate model to study TDP1 function. We combine knockdown of *sprtn* with a Tdp1-deficient zebrafish model and RPE1 cell lines with reduced expression of *TDP1* and *SPRTN*. For this purpose, we developed and verified new set of tools, including a Tdp1-deficient zebrafish strain, a zebrafish Tdp1 antibody, morpholino probes for *sprtn* silencing in embryos, a modification of the RADAR (Rapid Approach to DNA Adduct Recovery) assay for DPC isolation from tissues, and detection of TOP1- and H3-DPCs in embryonic tissues. We show that TDP1 and SPRTN are required for *in vivo* resolution of TOP1 and H3 DPCs and that both enzymes are also required for repair of other cellular DPCs. In contrast to H3-DPC repair, where SPRTN and TDP1 function together, we show that they work in separate pathways during repair of endogenous TOP1-DPCs. Our results reveal the relationship between TDP1 and SPRTN in the repair of DNA-protein crosslinks (DPCs) in human cells and in animal model, providing the first in-depth insights into the repair of DPCs at the organism level.

## Materials and methods

### Zebrafish husbandry and chemical treatment of embryos

Zebrafish (*Danio rerio*) AB were purchased from the European Zebrafish Resource Centre (EZRC, Karlsruhe, Germany). Fish were maintained at a constant temperature of 28 ᵒC and on a 14-hour light and 10-hour dark cycle, with water quality (temperature, pH, and conductivity) monitored daily. Embryos were maintained in E3 media (5 mM NaCl, 0.17 mM KCL, 0.33 mM CaCl_2_, and 0.33 mM MgSO_4_) in petri dishes in the incubator at 28 °C until 2dpf. Two-day-old embryos were dechorionated and treated with 10 µM camptothecin (CPT, Alpha Aesar: J62523) for 1 hour or 5 mM formaldehyde (FA, KEMIKA: 0633501) for 30 minutes. All handling and experiments were performed in accordance with the directions given in the EU Guide for the Care and Use of Laboratory Animals, Council Directive (86/609/EEC), and the Croatian Federal Act on the Protection of Animals (NN 135/06 and 37/13) under the project license HR-POK-023.

### Cell culture, siRNA and chemical treatments

HEK293T (Human embryonic kidney cells) and RPE1 (Retinal pigmentosum epithelial cells 1) cell lines were used in this study. Cells were cultured at 37°C in Dulbecco’s Modified Eagle’s medium (Capricorn, DMEM-HPA) containing 10% FBS (Capricorn, FBS-11A) serum under 5% CO_2_. RPE1 cells were seeded at 1×10^4^ cells/mL in 75 cm2 cell culture plates. The siRNAs targeting *TDP1*, *TDP2*, and *SPRTN* (Table 1) were transfected with Dharmafect reagent (Horizon Discovery, T-2005-01) according to the manufacturer’s instructions. Cells were incubated for 72 hours; they grew to 80-100% confluence and were collected for the RADAR assay and qPCR experiments. Silencing efficiency was determined by qPCR using the primers listed in Table 5. Before collection, cells were treated with 50 nM CPT for 1 hour in serum-free media or incubated with 1 mM FA for 20 minutes in ice-cold serum-free media.

**Table 1.**
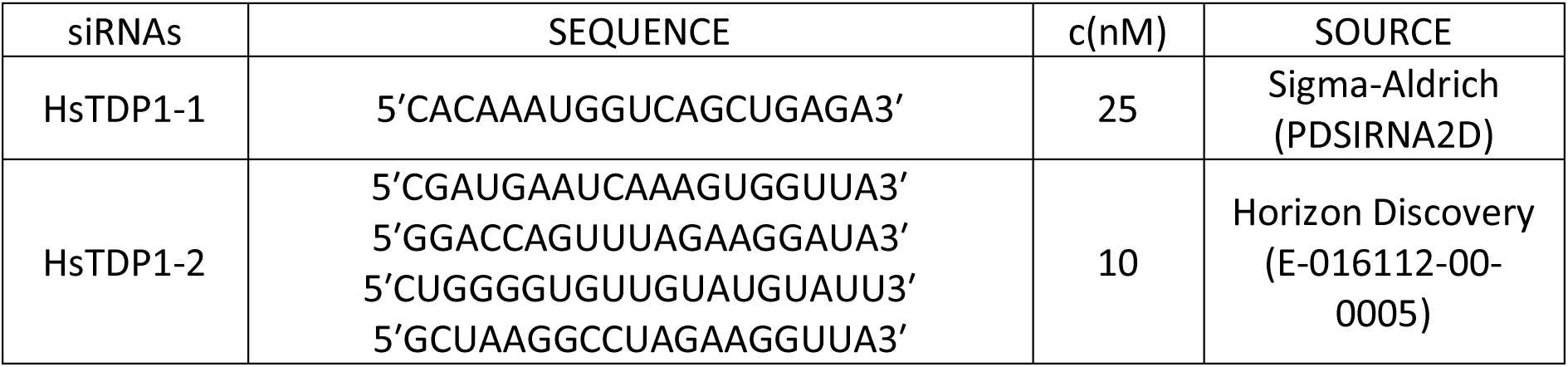
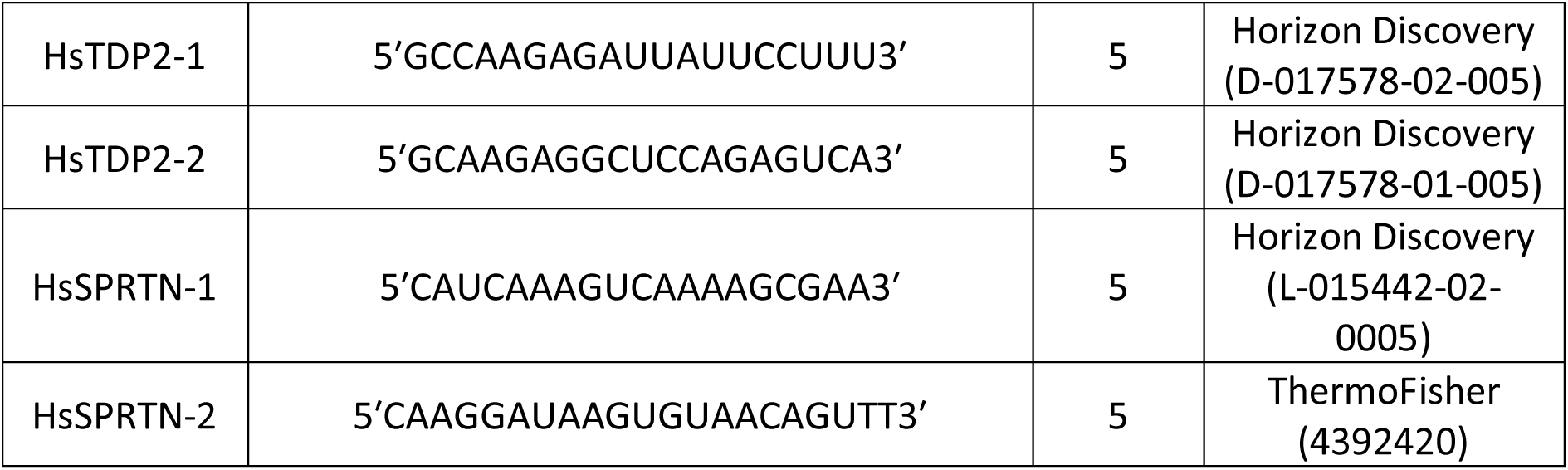
siRNAs used for gene silencing in RPE1 cells, with corresponding concentrations, manufacturers, and catalog numbers.

### Phylogenetic and syntenic analyses and structural modelling

Blast searches were performed using the NCBI database (National Center for Biotechnology Information) and human TDP1 sequence across bacteria, yeast, algae, plants, fungi, invertebrate and chordate species. Full length protein sequences were aligned using the MAFFT (Multiple Alignment using Fast Fourier Transform) alignment algorithm (Katoh et al., 2002). Alignment quality score was assessed using the Guidance2 server and was 0.563, indicating sufficiently high alignment quality for tree building (Penn et al., 2010). The phylogenetic tree was constructed using the Maximum Likelihood method (PhyML software with optimized tree topology, LG model, 8 rates of categories, tree searching operation best of NNI&SPR (Nearest Neighbor Interchange & Subtree Pruning and Regrafting) (Guindon & Gascuel, 2003). Branch support Alrt values (Approximate likelihood-ratio test) are shown at tree nodes on a scale of 0-1, where 1 is maximum node confidence (Anisimova & Gascuel, 2006). Synteny *tdp1* gene analysis between zebrafish, human and mouse were performed using Genomicus a browser for conserved synteny synchronized with genomes from the Ensembl database (Louis et al., 2013). Zebrafish Tdp1 was modeled using the Phyre2 workspace (Kelley et al., 2015) and human TDP1 (PDB: c1nopB) as a template. The degree of protein disorder was predicted using the PONDR-FIT software (Xue et al., 2010).

### Generation of the *tdp1* mutant zebrafish line

The short guide (sg)RNA (5′GAATGTGGGGGTCTCTTC3′) targeting exon 2 (Figure 1A) was selected using the CRISPR scan algorithm (Moreno-Mateos et al., 2015) and was generated as previously described (Modzelewski et al., 2018). A mixture containing 600 ng/uL of Cas9 protein (NEB: M0386) and sgRNA complex at a 1:1 ratio was generated with a total volume of 3 uL, and 1 nL was injected into one cell stage embryos. Three months after injection, when the F0 zebrafish reached adulthood, they were outcrossed to WT, and their progeny was analyzed using High-resolution melting analysis (HRMA) and sequencing to identify founder fish carrying the target mutation with the primers shown in Table 2. Founder fish were crossed and the F1 generation was raised and genotyped. Female and male were found to have a premature stop codon mutation (Figure S1E), and they were bred to produce the F2 generation harboring two changes in exon 2 of the *tdp1* gene.

**Figure 1.**
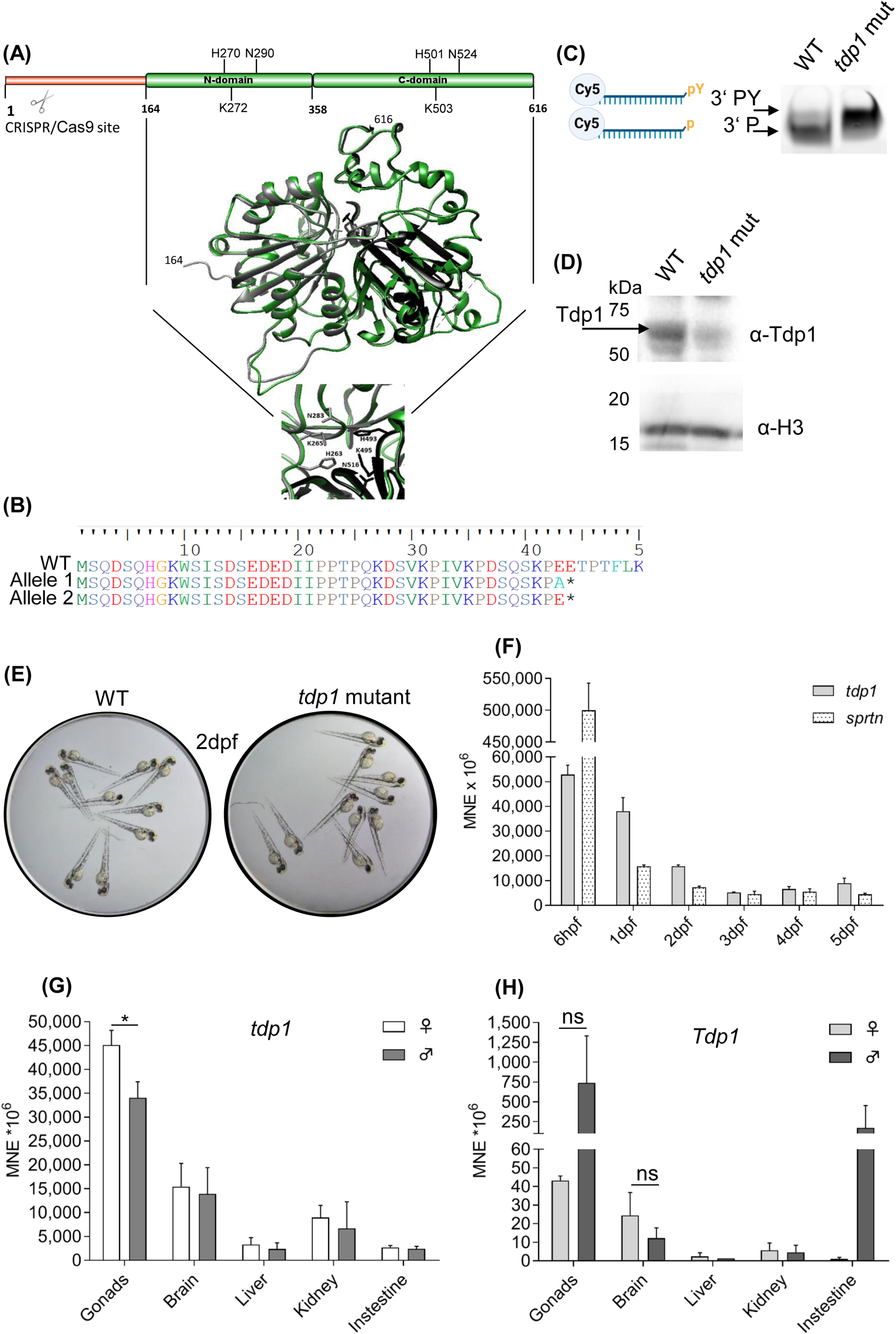
Structural comparison of zebrafish and human TDP1, validation and characterization of the zebrafish *tdp1* mutant line, and *tdp1* expression profiles in zebrafish embryos and zebrafish and mouse tissues. A) The zebrafish Tdp1 structural model (in green) is overlapped with the human TDP1 crystal structure (PDB: 1jy1) (Davies et al., 2002), shown in grey (N domain) and black (C domain). Zebrafish Tdp1 was modelled using the Phyre2 workspace (Kelley et al., 2015) according to the human TDP1 (PDB: c1nopB). N domain and C domain form a pseudo-2-fold axis of symmetry where each domain contributes to the active site: H263, K265 and N283 in the N domain and H493, K495 and N516 in the C domain. B) Amino acid sequence of Tdp1 in *tdp1* mutant fish line: frameshift and introduction of a premature stop codon in *tdp1* mutant fish line is deduced from DNA sequencing (*, premature STOP). C) TDP1 activity assay performed with 600 ng of lysate from 2-dpf WT and *tdp1* mutant embryos. Left panel: scheme created with BioRender.com of TDP1 substrate oligonucleotide with tyrosine (pY) on 3’ end and Cy5 fluorescent reporter on 5’ end and a reaction product after TDP1-mediated removal of tyrosine (p); right panel: TDP1 activity assay reactions resolved on 20% homemade urea gel and visualized using the ChemiDoc MP Imaging System to detect Cy5 fluorescence D) Western blot using a custom antibody against zebrafish Tdp1 shows the absence of a specific Tdp1 signal (68 kDa, indicated by arrow) in *tdp1* mutant embryo lysate. Histone H3 was used as a loading control. E) Images of WT and *tdp1* mutant embryos (2dpf, 2 days post fertilization). Embryos were maintained in E3 media, placed on a lid of a 96-well culture plate, and visualized with stereo microscope (Motic-SMZ-171-TP). Images were captured using a Canon 250D DSLR camera. F) *Tdp1* and *sprtn* expression patterns during the embryonic development from 6 hours post fertilization (6 hpf) to 5 days post fertilization (5 dpf). Data represent MNE (mean normalized expression) ± SD (n = 3) normalized to the housekeeping gene *atp50*. G) Tissue expression pattern of *tdp1* in male and female zebrafish, with statistically significant differences between expression in ovaries and testes (\**p* < 0.05) determined by unpaired t-test. Data are presented as MNE (mean normalized expression) ± SD (n = 3) normalized to the housekeeping gene *atp50.* H) Tissue expression pattern of *Tdp1* in male and female mice (ns, non-significant, *p* > 0.05). Data represents MNE (mean normalized expression) ± SD (n = 3) normalized to the housekeeping gene *Atp50*.

**Table 2.**
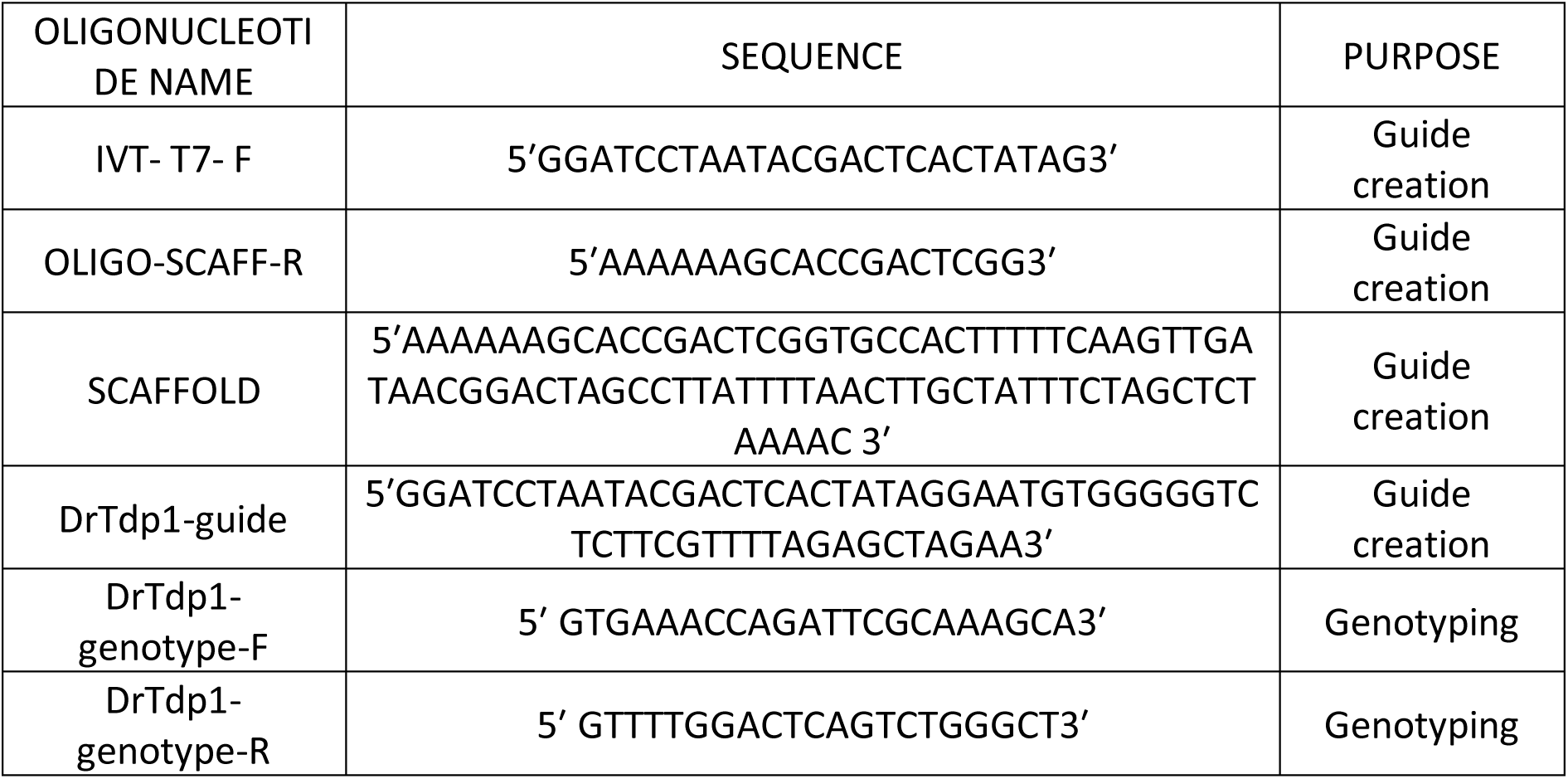
List of oligonucleotides used for the tdp1 zebrafish mutant line creation and genotyping. All oligos were purchased from Macrogen (Europe).

### sgRNA synthesis and microinjecting procedure

To generate a *tdp1* mutant line, we targeted the first exons of zebrafish Tdp1, and the CRISPR scanning algorithm selected (5’GAATGTGGGGGTCTCTTC3’) as the sgRNA with the highest score of 54 and a low off-target effect of 4.63 CFD (Moreno-Mateos et al., 2015). The gRNA was generated according to the established protocol of (Modzelewski et al., 2018). Briefly, the sgRNA template was inserted between two short DNA sequences, one containing the T7 promoter and the second sequence complementary to the scaffold template. In the PCR mixture, we combined the guide DNA template for the sgRNA, the IVT_ T7_ F primer that anneals at the beginning of the template (T7 promoter), the scaffold template, and the OLIGO_SCAFF_R primer that sits at the end of the scaffold sequence (Table 2). The PCR product serves as the template for *in vitro* T7 transcription using the MEGAshortscript™ T7 Transcription Kit (Invitrogen™, AM1354). *In vitro* transcription was performed according to the manufacturer’s instructions, and the transcribed RNA was purified using the Monarch® RNA Cleanup Kit (NEB, T2040L), RNA concentration was measured, and the guide RNA was stored at -80 °C. On the day of injection, guide RNA was incubated with Cas9 protein in a 1:1 ratio for 5 min at 37 °C. The sgRNA/Cas9 complex was pipetted into a Femtotips needle (Eppendorf, 5242957000) and connected to a microinjector (Eppendorf, FemtoJet® 4x).

### Tdp1 activity assay

The activity assay was performed as described in (Zaksauskaite et al., 2021). Briefly, embryo lysates were incubated with a TDP1 oligonucleotide substrate (5′GATCTAAAAGACT3′) containing a tyrosine at the 3’ DNA end and Cy5 at the 5’ end for visualization, which was purchased from Midland Certified Reagent Company. If TDP1 is active, a shift in the size of the oligonucleotide substrate becomes visible, revealing the removal of the tyrosine from the substrate. In brief, two-day-old mutants and WT embryos were deyolked in ice-cold PBS, and pelleted embryos were homogenized twice for 10 seconds (Ultra Turrax T25, IKA - Janke & Kunkel) in 200 µL of lysis buffer (200 mM Hepes, 40 mM NaCl, 2 mM MgCl2, 0.5% Triton X-100 with proteinase inhibitors (leupeptin, aprotinin, chymostatin, pepstatin at a concentration of 1 µg/mL and PMSF at a concentration of 1 mM)) and incubated on ice for 30 minutes. The supernatant protein solution (600 ng) was incubated with 2.5 µM labeled oligonucleotide substrate (Midland Certified Reagent Company) in assay activity buffer (25 mM Hepes (pH 8.0), 130 mM KCl, and 1 mM dithiothreitol (DTT)) to a final reaction mixture of 10 uL. The reaction was incubated at 37 ⁰C for 1 hour, then loading buffer was added, followed by boiling at 90 ⁰C for 10 minutes. All samples were loaded onto a pre-run 20% homemade urea gel and run at constant voltage (150 V) for approximately 1 hour. The oligonucleotide substrate products were visualized using the ChemiDoc MP Imaging System to detect Cy 5 fluorescence (Bio-Rad, 1708280).

### Zebrafish gene silencing with morpholino oligonucleotides

Two morpholino oligonucleotides, one targeting the 5’-UTR region of the zebrafish *sprtn* gene and the other targeting the exon 2-intron 2 boundary were designed to block zebrafish *sprtn* transcription, ordered from Genetools (USA), and used as described in (Nasevicius & Ekker, 2000) (Table 3). 1 nL of the microinjection mixture containing 250 µM of each MO, 0.3 M KCl and 0.015% phenol red was injected into each one-cell stage embryo. Two-day-old morphants were collected as follows: i) 5 embryos for qPCR analysis and to check the efficiency of morpholino splice blocking, ii) 30 embryos for RADAR experiments. Total RNA was extracted using the Monarch Total RNA Kit (NEB, T2040) according to the manufacturer’s instructions. Equal amounts of RNA were reverse transcribed using the High-Capacity cDNA Kit (Applied Biosystems™, 4368814). The splice-blocking efficiency of morpholino was verified by PCR on 5 ng of cDNA with primer pairs annealing upstream and downstream of the morpholino target (Table 4): the 439 bp amplicon from WT and the 339 bp amplicon from *sprtn* morphants caused by exon 2 skipping were separated by 1% agarose gel electrophoresis. Silencing efficiency was quantified using Image J (Figure S2A and B). The same cDNA was used for qPCR analysis with the primers listed in Table 5.

**Table 3.**
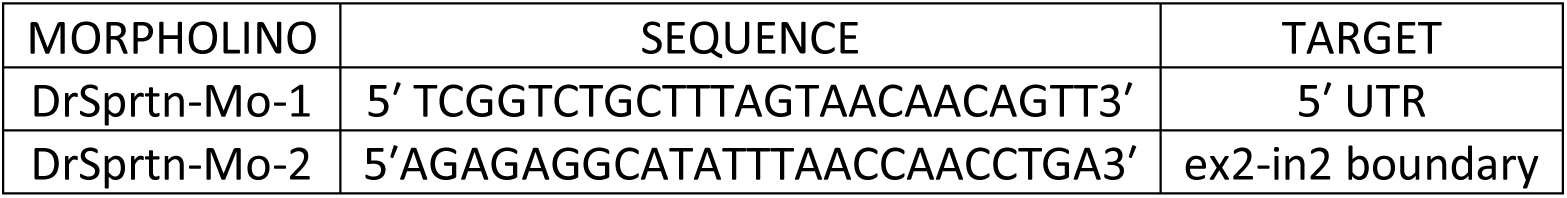
List of morpholinos used to silence *sprtn* expression in zebrafish. The morpholinos were obtained from Genotools (USA).

**Table 4.**
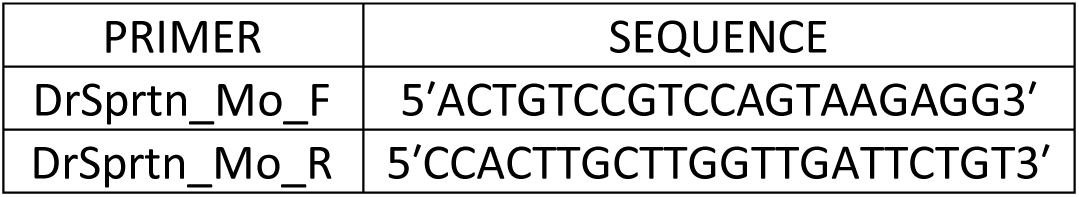
Primers used to determine the silencing efficiency of the exon skipping morpholino. The primers were obtained from Macrogen (Europe).

### DPC isolation, detection, and quantification

DPC isolation and detection was performed using the modified RADAR (rapid approach to DNA adduct recovery) assay (Kiianitsa & Maizels, 2013). We modified RADAR assay in order to increase processivity and reduce variability between experiments. We also optimized DPC isolation from zebrafish embryos (1-3 dpf). The main modification included the use of a lyophilizer which substituted TCA precipitation step, thus significantly reducing sample loss and variability between samples. DPC isolation from cells was performed with 8 x 10^6^ cells grown in a 75 cm^2^ cell flask. DPC isolation from 2 dpf embryos was performed as follows: (1) lysis of 30 embryos per condition with pre-warmed lysis buffer (6 M GTC, 10 mM Tris-HCl (pH 6.), 20 mM EDTA, 4% Triton X100, 1% N-lauroylsarcosine sodium, and 1% β-mercaptoethanol) for 10 min at 50 ᵒC; (2) precipitation of DNA with crosslinked proteins by adding an equal volume of 98% ethanol; (3) centrifugation at 10,000 rcf for 10 min at 4 ᵒC; (4) washing the pellet four times with wash buffer (20 mM Tris (pH 7.4), 1mM EDTA, 50 mM NaCl, 50% EtOH) with centrifugation steps at 10,000 rcf at 4 ᵒC; (4) dissolving the pellet in 8 mM NaOH. A 25 µL aliquots of the DPC sample was treated with proteinase K (15 µg, Fisher Scientific, BP1700-100), and the DNA was quantified using Pico Green according to the manufacturer’s instructions (Invitrogen, P7581). The DPC samples were normalized to the one with the lowest amount of DNA and treated with DNAse (Millipore, E1014) for 1 h at 37 °C. They were then snap-frozen in liquid nitrogen, submitted to overnight lyophilization, and then dissolved in SDS loading buffer (4 M urea, 62.5 mM Tris-HCl (pH 6.8), 1 mM EDTA, 2% SDS). Total DPCs were resolved by SDS-PAGE gradient gel (5-18%) and detected by silver staining according to the manufacturer’s protocol (Sigma Aldrich, PROTSIL1). For specific DPC detection, DPCs isolated from cells were applied to the nitrocellulose membrane (GE Healthcare, 10-6000-02) for dot blot analysis, while DPCs isolated from embryos were first resolved by SDS-acrylamide gradient gel (5-18%), transferred to PDF membrane (Roche, 03010040001) and then visualized by Western blotting. To detect the presence of histone H3 or TOP1 in the DPCs, the membranes were immunostained with the specific antibodies. For detection of histone H3, 200 ng of DNA-normalized DPCs were subjected to dot blot or Western blotting, and then immunostained with anti-H3 antibody (Cell Signaling, #9715, 1:3000). For detection of TOP1, DPC equivalent of 500 ng DNA were applied to a dot blot (Bethyl, A302-589A, 1: 1000), and an equivalent of 1 µg of DNA-normalized DPCs was resolved using SDS-acrylamide gradient gel with the addition of 4 M urea (Cell Signaling, #38650, 1: 1000). DNA detection was performed by applying 2 ng of the sample collected for Pico Green detection to a nylon membrane and detecting with the α-dsDNA antibody (abcam ab27156, 1:7000).

### MTT viability test

The MTT colorimetric assay was used as an indicator of cell viability as previously described (Wang et al., 2010). In brief, RPE1 cells were seeded in 24-well plates and transfected with siRNA. After 72 hours, cells were incubated with 100 μl of a 5 mg/mL MTT solution (Alfa Aesar, L11939) for 3 hours. The solution was removed and 500 μL isopropanol (Kemika, 1622601) was added, followed by shaking at 350 rpm for 15 minutes (BioSan, PST -60HL-4, Plate Shaker-Thermostat). Absorbance was measured at 570 nm using a microplate reader (Tecan, Infinite M200).

### Quantitative PCR analysis

qPCR analysis was performed with the GoTAQ qPCR mix (PROMEGA, A6001) using primer pairs that span exon-exon boundaries to exclude possible amplification from genomic DNA (Table 5). Target gene expression was normalized to a housekeeping gene, the *ATP50* gene (Otten et al., 2018), which is expressed equally in all samples (embryonic developmental stages and adult tissues, data not shown). Quantification was performed using the Q-gene (Muller et al., 2002) method and gene expression is expressed as MNE (Mean normalized Expression) where ***MNE = E*(*HKG*)^*Ct*(*HKG*)^/*E*(gene)^*Ct*(*gene*)^ × 10^6^**. E (HGK) is the primer efficiency of the housekeeping gene; E (gene) is the primer efficiency of the target gene; Ct (HKG) is the mean Ct value for the housekeeping gene; and C t (gene) is the mean Ct value of the target gene. The expression level of the target gene was then normalized to the housekeeping gene *ATP50*.

**Table 5.**
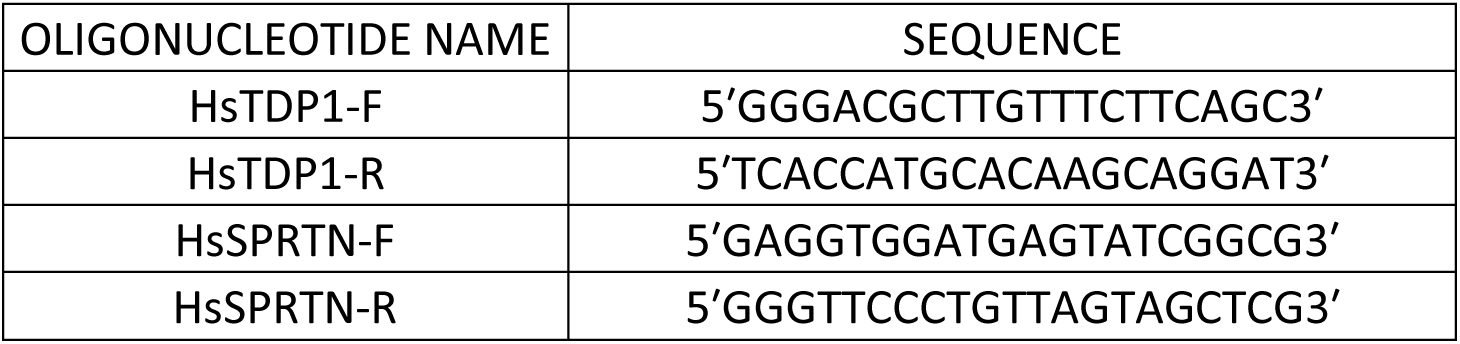
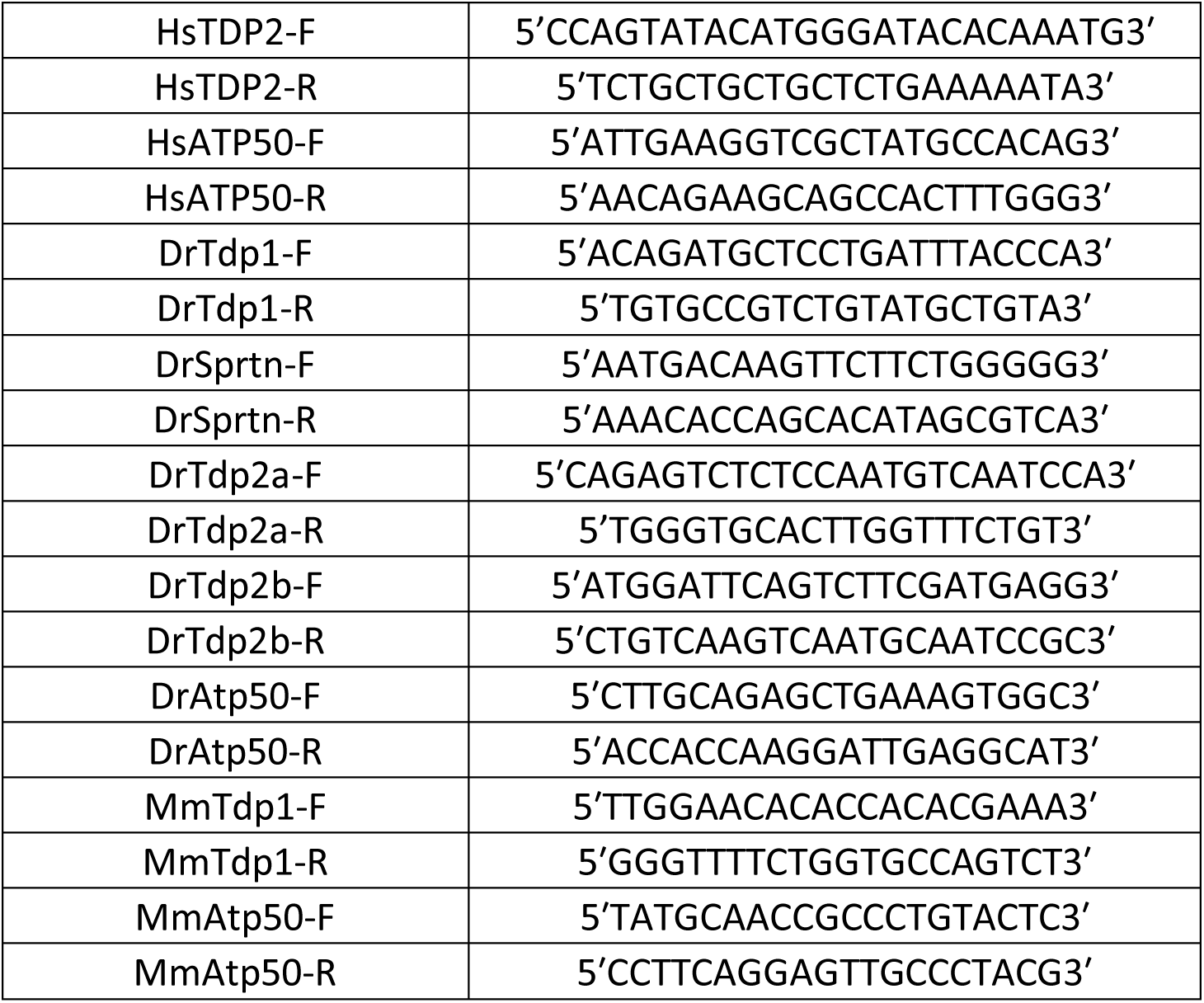
List of primers used for human (Hs), zebrafish (Dr), and mouse (Mm) qPCR analysis. Primers were purchased from Macrogen (Europe).

### High-resolution melting curve (HRM) analysis

High-resolution melting curve analysis was used to distinguish mutants from WT fish with differences in dsDNA melting fluorescence. Primer par of Table 2, genomic DNA, MeltDoctor HRM Mastermix (Applied Biosystems™, 4415440) were added to 10 µL PCR mix and fluorescence was detected using StepOnePlus™ Real-Time PCR System (Applied Biosystems™, 4376600). Data were analyzed using High Resolution Melting (HRM) software and melting curve changes were confirmed by sequencing.

### Development and verification of a custom-made zebrafish Tdp1 antibody

The peptide N-GALEKNNTQIMVRSYE-C was specifically chosen to detect both zebrafish and human TDP1 proteins and was used to immunize two rabbits followed by affinity purification of serum (Genosphere, UK). To test the specificity of the antibody, we performed western blotting on WT and *tdp1* mutant zebrafish samples, as well as on WT HEK293T cells and HEK293T cells overexpressing human TDP1. Cells were transfected with the recombinant plasmid carrying the human *TDP1* gene (GenScript, OHU22350D) using PEI transfection reagent as previously described (Popovic et al., 2013; Tom et al., 2008). Cells were collected after 72 hours and lysed in RIPA buffer (150 mM NaCl, 1% Triton X-100, 0.50% Na-deoxycholate, 0.10% SDS, 50 mM TrisHCl (pH 8)) followed by a 10-second sonication. 2dpf zebrafish embryos were lysed in RIPA buffer and sonicated twice for 10 seconds. 30 µg of protein extract from cells and 50 µg from zebrafish embryos were loaded on an SDS acrylamide gel, transferred to a PVDF membrane, blocked with milk for 2 hours, and incubated overnight with the custom-made anti-Tdp1 antibody (1:1000). The blot was visualized by incubating the membrane with an HRP-labelled anti-rabbit antibody followed by detection using ECL (Biorad) at the Chemidoc (Biorad).

### Western blotting and Dot blotting

The cell/embryo lysates and isolated DPCs were mixed with 5x Laemmli buffer containing 50 mM Tris-HCl (pH 6.8), 2% SDS, 10% w/v glycerol, 0.05% bromophenol blue, and 5% β-mercaptoethanol. Additionally, the cell or embryo lysates were boiled for 10 minutes at 90° C. The proteins were then separated by 5-18% SDS-PAGE gradient gels and transferred to PVDF membranes. Specific DPCs in cells were detected using a dot blot (Bio-Rad, Bio-Dot Apparatus 1706545) and transferred directly to a nitrocellulose membrane by vacuum aspiration. The membranes were blocked with 5% low-fat milk (Roth, T145.1) in TBST buffer (10 mM Tris-HCl (pH 7.5), 15 mM NaCl, 0.02% Tween 20) for 2 hours at room temperature and then incubated with the corresponding primary antibody in 2.5% BSA in TBST overnight at 4°C. The membranes were then incubated with an HRP-labelled anti-rabbit antibody (Sigma-Aldrich. a0545, 1:100000) for 1 hour and washed three times for 15 minutes in TBST. Detection was performed according to the manufacturer’s instructions for the ECL blotting substrate (Bio-Rad, 1705061) and visualized using the ChemiDoc™ XRS+ System (Bio-Rad, 1708299).

### Statistical Analysis

The ImageJ software (Abràmoff et al., 2004) was used to quantify dot blots, Western blots, and morpholino-mediated silencing efficiency in zebrafish. Graphical representation of the expression data and statistical analysis was conducted using the unpaired two-sided Student’s t-test using GraphPad Prism 8. Differences between two independent variables were considered significant when p < 0.05. All experiments were performed three to six times following the setup for biological replicates for RADAR isolation from cells and embryos, with means ± SD shown in each column.

## Results

### Comparison of zebrafish, mouse and human TDP1 proteins: phylogeny, synteny, sequence and structure

We constructed a phylogenetic tree of Tdp1 orthologs in multicellular organisms, yeast, and bacteria by aligning protein sequences using the MAFFT algorithm (Katoh et al., 2002) and building a phylogenetic tree using Maximum Likelihood method (Guindon & Gascuel, 2003). The Tdp1 protein is very conserved in all ‘kingdoms’ of life, from bacteria and yeast to plants and animals and is always present as a single ortholog (Figures S1A and S1B and Table S1). Interestingly, the Tdp1 orthologs in invertebrates form two distinct clusters: the one which is phylogenetically very close to the vertebrate cluster and the other one which is closer to the yeast and bacterial orthologs (Figures S1A and S1B). Human and zebrafish Tdp1 are phylogenetically very close (Figures S1A) and structurally very similar (Figure 1A). We modeled the structure of zebrafish Tdp1 using the crystal structure of human TDP1 (PDB: c1nopB) (Davies et al., 2002) in the Phyre2 workspace (Kelley et al., 2015). These orthologs share a very similar structure of N and C domains with a remarkable degree of conservation (Figure 1A). N domain (164. – 358. amino acid) and C domain (359. – 616. amino acid) form a pseudo-2-fold axis of symmetry with each domain contributing histidine, lysine and asparagine to the active site: H263, K265, and N283 in the N domain and H493, K495, and N516 in the C domain (Davies et al., 2002; Flett et al., 2018) (Figure 1A). Upstream of the N domain, is an N terminal portion which is heavily disordered in both zebrafish and human TDP1 (1-163. and 1-144. amino acids, respectively) (Figure S1C). This part is highly variable among species, and its structure has not yet been solved. The amino acid sequence similarity between human and zebrafish Tdp1 is 66% (identity 55%), whereas the similarity between mouse and human is 83% (identity 77%). If we exclude the variable N terminus, similarities between orthologs are much higher: 76% between human and zebrafish (identity 66%) and 92% between human and mouse (identity 88%). After determining the phylogenetic, structural, and sequence similarities between the human, mouse, and zebrafish Tdp1 proteins, we analysed the gene environment of the orthologs and found that it is partially conserved. Zebrafish *tdp1* on chromosome 17 is surrounded by the upstream neighboring gene *kcnk13a* and the downstream neighboring genes *efcab1*1 and *foxn3* similarly as in human and mouse *TDP1*genes located on chromosomes 14 and 12, respectively (Figure S1D). Apart from the aforementioned neighboring genes, the gene environment between zebrafish on one side and human and mouse *TDP1* on the other side is not conserved. In contrast, the gene environment of *TDP1* in humans and mice exhibits preserved genomic order that was presumably passed down from a common mammalian ancestor (Figure S1D).

### Generation and characterization of zebrafish line lacking Tdp1 protein

The CRISPR/Cas9 system was used to generate a zebrafish strain lacking the Tdp1 protein. A gRNA targeting exon 2 induced a frameshift mutation that resulted in a premature stop codon at amino acid position 44 upstream of catalytic residues H270, K272, and N290 in the N domain (Figures 1A and 1B). The gRNA/Cas9 complex was injected into one-cell stage embryos and fish were grown to adulthood. Identified founder fish in the F0 generation that produced germlines with a frameshift mutation that resulted in premature stop at amino acid position 44 were crossed and the F1 generation was raised. When the F1 generation reached adulthood, individuals were genotyped based on fin tissue and allele changes were sequenced. Female and male carrying described frameshift mutations (Figures 1B and S1E) were further crossed to produce an F2 generation deficient in Tdp1 protein (Figure 1B). The lack of functional Tdp1 was confirmed by enzyme activity assay (Figure 1C) and Western blot using a custom-designed antibody for zebrafish Tdp1 (Figures 1D and S1F). For the activity assay, we used a model substrate of Tdp1, a 3’-phosphotyrosyl-DNA probe (3’pY) with a fluorescent reporter Cy-5 at the 5’ end. When Tdp1 is active, tyrosine is removed from the substrate, resulting in the oligonucleotide form (3’p) (Figure 1C). Lysates from WT embryos (with active TDP1) were incubated with the labeled substrate and very efficient tyrosine removal was observed, whereas no reaction occurred after incubation with the lysates from *tdp1* mutant embryos demonstrating the absence of Tdp1 in the mutants (Figure 1C). Using the custom-made Tdp1 antibody, we showed that the Tdp1 protein is indeed absent in the *tdp1* mutant line (Figure 1D). The zebrafish antibody was intentionally designed with an epitope in a conserved protein region that overlaps between zebrafish and human TDP1 so that it could also be used to detect TDP1 in human cells. Thus, we were able to test the specificity of the new antibody by overexpressing human TDP1 in HEK293 cells (Figure S1F). After determining that the Tdp1 protein was indeed absent in the *tdp1* mutant line (Figure 1D), we examined whether Tdp1 deficiency resulted in phenotype changes in embryos and adult zebrafish. There were no obvious morphological differences between WT and *tdp1* mutant embryos, nor in adult zebrafish that are now 8 months old (Figures 1E and S1G). Future studies are needed to investigate specific phenotypes in adult Tdp1-deficient zebrafish, particularly with regard to neurodegeneration in old fish.

### Tdp1 is highly expressed throughout the embryonic development and in adult tissues

WT embryos were collected at different time points, starting at 6 hours post fertilization (hpf), when most of the maternal transcriptome is degraded (Mathavan et al., 2005). We show that *tdp1* is strongly expressed throughout embryonic development from 6 hpf to 5 days postfertilization (dpf) (Figure 1F). Expression levels are highest at early stages, at 6 hpf and 1 dpf, followed by a 5-fold decrease at later stages (2-5 dpf). Overall, the expression levels remain high from 6 hpf to 5 dpf (Figure 1F). To facilitate comparison of expression levels, we set arbitrary thresholds following previous publication (Loncar et al., 2016): high expression when MNE is > 60 x 10^6^ (Ct values > 22), moderate when MNE is 2 x 10^6^ – 60 x 10^6^ (Ct = 23 - 26), and low when MNE is < 2 x 10^6^ (Ct > 27). *Sprtn* shows a similar expression pattern to *tdp1* at later stages, with high and stable expression levels at 2 to 5 dpf (Figure 1F). *Sprtn* expression is particularly high at the 6-hpf, when mRNA levels are 33-fold higher than those of *tdp1*. In adults, *tdp1* is highly expressed in all analysed tissues, with the highest expression in testis and ovary, followed by a 3.3-fold lower expression in brain and kidney and a 14.7-fold lower expression in liver and intestine (Figure 1G). Gender differences are not significant except in the gonads, where *tdp1* is expressed 1.3-fold more in the ovaries than in the testes.

To compare *tdp1* expression in zebrafish with the most commonly used animal model, the laboratory mouse, *Tdp1* expression was measured in the corresponding mouse tissues (Figure 1H). The expression pattern is generally similar to that of the zebrafish with the highest expression in the gonads, followed by the brain, liver, kidney, and female intestine. However, there are some differences between the two animal models: (1) overall expression levels are much lower in mice, with moderate expression in testis and low expression in all other tissues, in contrast to the high expression in all zebrafish tissues; (2) in mice, *Tdp1* is expressed 17.2-fold less in ovaries than in testes, in contrast to 1.3-fold higher expression in zebrafish ovaries compared with testes; and finally, (3) a gender difference in *Tdp1* expression in mouse intestine was not observed in zebrafish (Figure 1H).

### Tdp1 repairs Top1- and histone H3-DPCs *in vivo*

DPC isolates were analysed for the presence of TOP1- and histone H3-DPCs by Western blot and dot-blot using protein-specific antibodies. Four biological replicates were used for detection of Top1- and H3-DPCs in zebrafish embryos, whereas three biological replicates were used for detection in RPE1 cells. Tdp1-deficient embryos exhibit greatly increased Top1-DPC levels under physiological conditions (4.8-fold more than WT) (Figures 2A and 2B). In comparison, the effect of the Top1-DPC inducer, CPT (10 µM, 1 hour) was weaker in WT embryos: 2.6-fold increase compared to WT (Figures 2A and 2B). CPT further increased Top1-DPC levels in *tdp1* mutants (6.2-fold) (Figures 2A and 2B). FA treatment (5 mM, 30 min) had a much stronger effect on Top1-DPC induction than CPT in both, WT, and mutant embryos: FA induced Top1-DPC levels by 7.1-fold in WT and by 9-fold in *tdp1* mutant embryos (Figures 2A and 2B).

**Figure 2.**
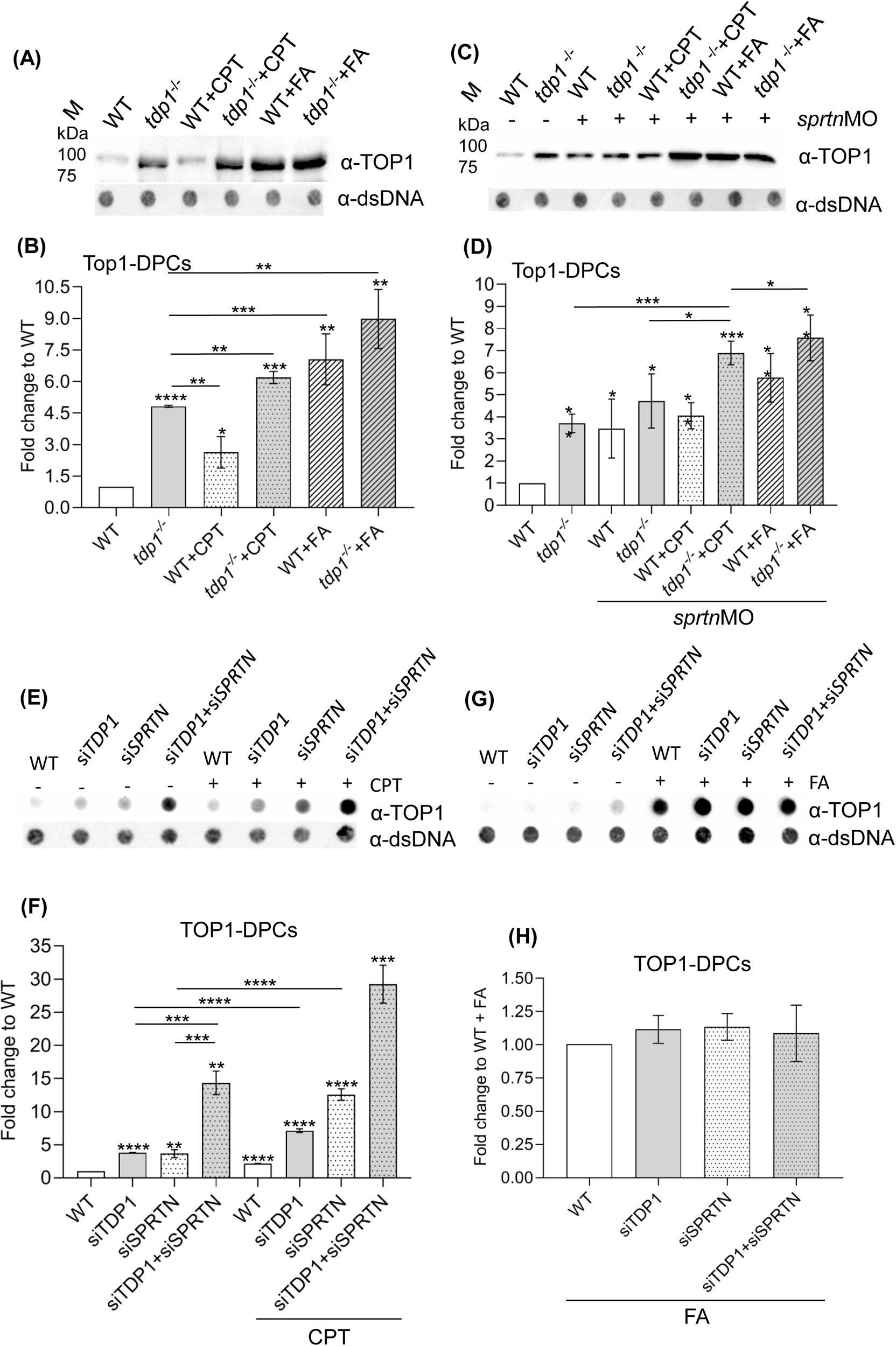
Tdp1 deficiency causes strong accumulation of endogenous and chemically induced Top1-DPCs in embryos and cells. (A) Western blot showing zebrafish Top1-DPCs in *tdp1* mutant embryos before and after camptothecin (CPT) (10 µM, 1 hour) and formaldehyde (FA) treatment (5 mM, 30 min) (DPC equivalent of 1 µg DNA was loaded per well) and (B) corresponding quantification (n = 4). (C) Western blot showing zebrafish Top-1 DPCs in *tdp1* mutant embryos before and after *sprtn* silencing and CPT (10 µM, 1 hour) and FA treatment (5 mM, 30 min) and (D) corresponding quantification. E) Dot blots showing human TOP1-DPCs detected with TOP1-specific antibody before and after CPT treatment of RPE1 cells (50 nM, 1 hour) with corresponding DNA loading controls (DPC equivalent of 500 ng DNA was loaded per well). F) Quantification of E from three different biological replicates normalized to untreated WT cells. G) Dot blots showing human TOP1-DPCs detected with TOP1-specific antibody before and after FA treatment of RPE1 cells (1 mM, 20 min) with corresponding DNA loading controls (DPC equivalent of 500 ng DNA was loaded per well). H) Quantification of G (n = 3). Results represent mean fold change ± SD of three different experiments. Statistically significant changes as a result from unpaired Student’s t-test are shown as * (*p* < 0.05), **(*p* < 0.01), *** (*p* < 0.001) or **** (*p* < 0.0001).

In RPE1 cells, the pattern of TOP1-DPC induction (Figure 2E-H) was to some extent similar to that in embryos. TOP1-DPCs strongly accumulated in RPE1 cells after *TDP1* silencing with 3.7-fold increase compared to endogenous levels (Figures 2E and 2F, Figure S2C). In comparison, CPT, a TOP1-DPC inducer, caused a 2.2-fold increase in WT cells (Figures 2E and 2F). When cells were further challenged by the combination of TDP1 deficiency and CPT treatment, TOP1-DPC levels increased 7.1-fold compared to nontreated WT cells (Figures 2E and 2F), confirming that TDP1 is critical for TOP1-DPC removal in human cells. In comparison, FA treatment (1 mM, 20 min) had a very strong effect on TOP1-DPC increase in WT cells, and this induction did not further increase after *TDP1* or/and *SPRTN* silencing (Figures 2G and 2H).

To investigate whether TDP1 is involved in the repair of DPCs other than TOP1-DPCs, we chose to examine histones as possible TDP1 substrates based on the recent *in vitro* data of Wei et al. (2022), who showed that purified TDP1 removes crosslinked H4 and H2B at abasic sites. We chose histone H3 as a representative of the core histones because we had optimized protocols and a very sensitive antibody for its detection in the DPC isolates. In the embryos, Tdp1 deficiency caused very strong accumulation of endogenous H3-DPCs: a 4.2-fold increase compared with WT embryos (Figures 3A and 3B). This is a very similar pattern to the one observed for the canonical substrate of Tdp1, Top1-DPC, in *tdp1* mutants (4.8-fold) (Figures 2A and 2B). CPT treatment caused a 4.7-fold increase of H3-DPCs in WT and even greater increase of 7.3-fold in *tdp1* mutants (Figures 3A and 3B), again consistent to the pattern of Top1-DPC accumulation after CPT exposure (Figures 2A and 2B). The FA treatment had a similarly strong effect in WT and *tdp1* mutant embryos, namely a 5.3- and 5.1-fold increase in H3-DPCs (Figures 3C and 3D).

**Figure 3.**
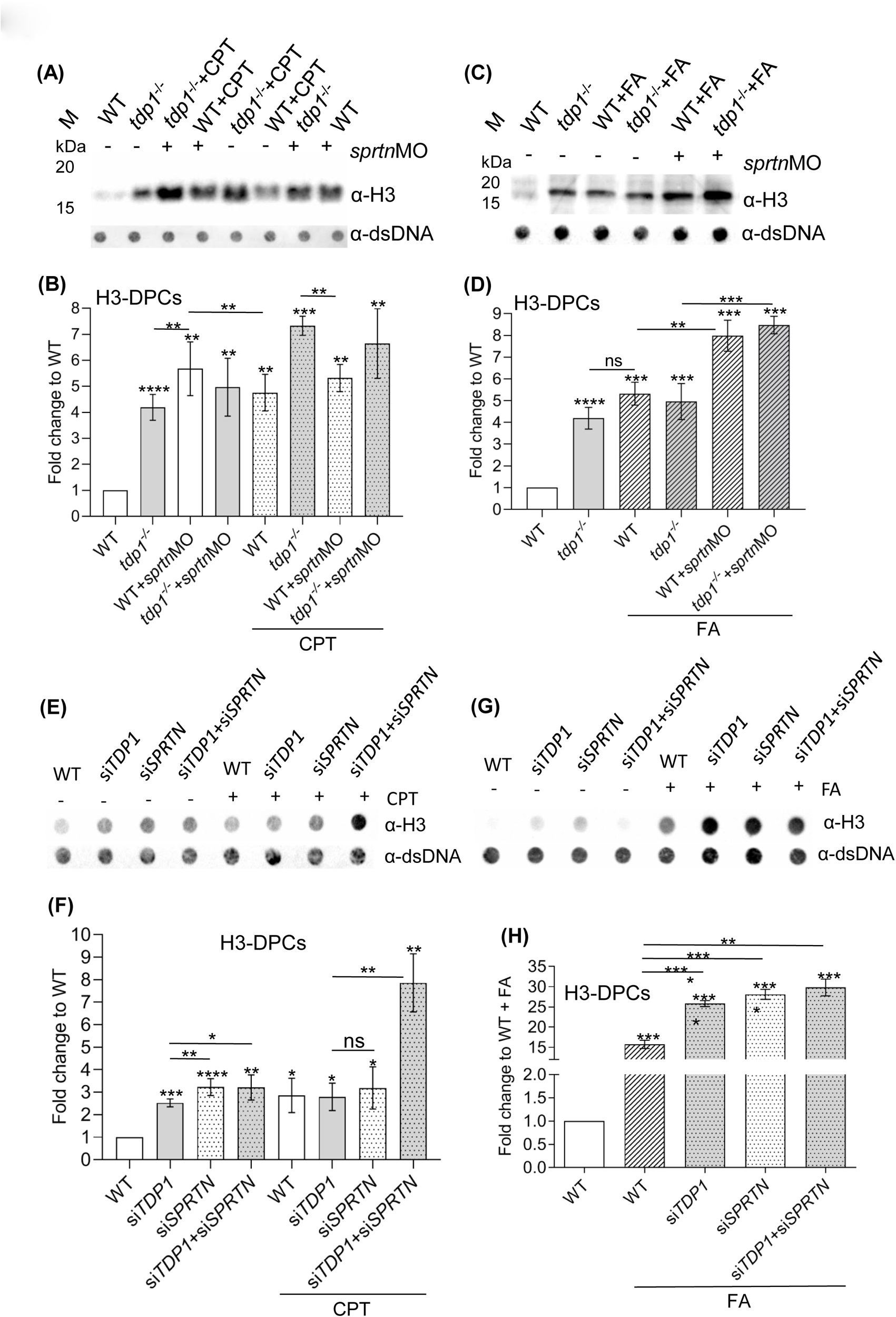
H3-DPC levels are increased *in vivo* in *tdp1* mutant fish line and in RPE1 cells with TDP1 deficiency. A) Western blot showing H3-DPC levels in *tdp1* mutant embryos in combination with *sprtn* knockdown and CPT (10 µM, 1h) treatment. Total DPCs were isolated from 2-day old embryos, separated by SDS-PAGE (DPC equivalent of 200 ng DNA per well) and detected with H3-specific antibody. B) Quantifications of H3-DPCs from four biological replicates with mean (± SD) fold change to endogenous H3-DPCs in WT embryos. C) Western blot analysis of H3-DPCs in zebrafish embryos and (D) corresponding quantification after FA treatment (5 mM for 30 min) (n = 4). E) Dot blots showing H3-DPCs after silencing *TDP1* and/or *SPRTN* before and after CPT exposure in RPE1 cells (50 nM CPT, 1h) and DNA loading controls. Equivalent of 200 ng DNA of total DPCs was loaded per sample. (F) Quantification of H3-DPC analysis in RPE1 cells (n = 3). G) Dot blots showing H3-DPCs after silencing *TDP1* and/or *SPRTN* before and after FA exposure (1 mM FA, 20 min) in RPE1 cells and DNA loading controls and H) Corresponding quantification (n = 3). Results are presented as mean ± SD with statistically significant changes, determined using an unpaired Student’s t-test, indicated with *(*p* < 0.05), **(*p* < 0.01), ***(*p* < 0.001), and ****(*p* < 0.0001).

In RPE1 cells, *TDP1* silencing induced H3-DPC levels by 2.5-fold compared with untreated WT cells (Figures 3E and 3F). It is important to note that CPT, although previously known to be a specific TOP1-DPC inducer, increased H3-DPCs 2.9-fold in WT cells (Figures 3E and 3F and S2E and S2F). H3-DPC levels were comparably affected when WT or *TDP1*-silenced cells were exposed to CPT: 2.9-fold and 2.8-fold increase, respectively (Figures 3E, 3F, S2E and S2F). In contrast, CPT induced many more H3-DPCs in *tdp1* mutants (7.3-fold), then in CPT-treated WT embryos (4.6-fold increase) (Figures 3A and 3B). FA caused a remarkable 15.7-fold increase in H3-DPCs in WT cells and a 25.8-fold increase in *TDP1*-silenced cells compared with WT untreated cells (Figures 3G and 3H).

### SPRTN proteolysis is necessary for TOP1 and H3 DPC repair *in vivo*

To test the hypothesis that upstream proteolysis by SPRTN is required for the subsequent action of TDP1 in removing TOP1-DPCs, we quantified TOP1-DPC levels in embryos and RPE1 cells under different conditions. In WT embryos, knockdown of *sprtn* using the morpholino approach reduced *sprtn* mRNA levels by 80% (Figures S2A and S2B) and caused a 3.5-fold increase in Top1-DPCs levels (Figures 2C, 2D and S2D). In *tdp1* mutant embryos, the increase in Top1-DPCs before and after *sprtn* knockdown was 3.5 and 4.7-fold, respectively (Figures 2C, 2D and S2D). Surprisingly, CPT treatment (10 µM, 1 hour) did not result in an additional increase in Top1-DPC levels in *sprtn-*silenced embryos (with functional Tdp1) (Figures 2C and 2D). Compared with untreated WT embryos, CPT exposure of embryos deficient in both Tdp1 and Sprtn increased Top1-DPC levels 6.9-fold which is a significant increase compared with *tdp1* mutants and *sprtn*-silenced mutants (Figures 2C and 2D). FA treatment (5 mM, 30 min) of *sprtn*-silenced embryos with functional Tdp1 caused a significant 5.8-fold increase in Top1-DPC levels compared with untreated WT embryos (Figures 2C and 2D) which is an additional increase in comparison to *sprtn*-silenced embryos (3.5-fold). In *sprtn*-silenced *tdp1* mutants, exposure to FA significantly increased Top1-DPC levels compared with nontreated *sprtn*-silenced *tdp1* mutants (p < 0.005) (Figures 2C and 2D).

In RPE1 cells, silencing of *SPRTN* with 65% efficiency (Figure S2C), resulted in a significant increase in TOP1-DPC levels (3.6-fold), which is a similar effect to silencing of *TDP1* (3.7-fold) (Figures 2E and 2F). When both *SPRTN* and *TDP1* were silenced (Figure S2C), TOP1-DPCs accumulated dramatically (14.3-fold increase), suggesting that both proteins are involved in TOP1-DPCs removal (Figures 2E and 2F). The same setup after CPT exposure (50 nM, 1h) showed a different pattern: *SPRTN* silencing caused a 12.6-fold increase, *TDP1* silencing 7.1-fold increase, whereas double silencing additionally increased TOP1-DPC levels by 29.2-fold (Figures 2E and 2F). FA treatment (1 mM, 20 min) dramatically increased TOP1-DPC levels to a similar extent under all conditions (Figures 2G and 2H).

In embryos, *sprtn* knockdown had a tremendous effect on H3-DPC levels in both, WT and *tdp1* mutants with 5.7- and 5.1-fold increase, respectively (Figures 3A and 3B). CPT (10 µM, 1 hour) caused a different pattern of H3-DPC induction in the same backgrounds: a 4.8-fold increase in *sprtn*-silenced embryos and a 6.6-fold increase in *sprtn*-silenced *tdp1* mutants (Figures 3A and 3B). Both inductions were weaker than in CPT-treated mutant embryos with functional Sprtn (7.3-fold) (Figures 3A and 3B). The levels of H3-DPCs observed in *tdp1* mutants after exposure to CPT show a significant increase compared to the endogenous levels of H3-DPCs in *tdp1* mutants (P < 0.0001). Moreover, this increase is even greater than the increase caused by *sprtn* silencing in *tdp1* mutants (P < 0.0001) (Figures 3A and 3B). At the same time, H3-DPC levels were similarly induced after *sprtn* silencing in WT and mutant embryos before and after CPT treatment (Figures 3A and 3B). Compared with CPT-treated WT or *tdp1* mutant embryos, knockdown of *sprtn* had no further effect on the increase in H3-DPCs (*p* < 0.05) (Figure 3A and 3B). *Sprtn* knockdown in FA-treated embryos (5 mM, 30 min) further increased H3-DPC levels compared in embryos with functional Sprtn: 8- and 8.5-fold increase in WT and mutants, respectively, versus 5.3- and 5.1-fold increases in WT and mutants with functional Sprtn, respectively (Figures 3C and 3D).

In RPE1 cells, SPRTN deficiency caused a very strong accumulation of H3-DPCs (3.2-fold increase) (Figures 3E, 3F, S2E and S2F). No additional effects on H3-DPC levels were observed when both *SPRTN* and *TDP1* were silenced. *SPRTN* silencing in untreated and in CPT-treated cells increased H3-DPCs similarly: by 3.2- and 3.2-fold, respectively (Figures 3E and 3F). However, simultaneous silencing of *TDP1* and *SPRTN*, followed by exposure to CPT had an additive effect on H3-DPC levels resulting in a 7.9-fold increase (Figures 3E, 3F, S2E and S2F). This increase is not as strong as for TOP1-DPC levels, where CPT treatment after simultaneous silencing dramatically increased TOP1-DPCs: from 14.3-fold to 29.2-fold (Figures 3E and 3F). When exposed to FA (1 mM, 20 min), *SPRTN*-silenced cells, as well as *SPRTN*- and *TDP1*-silenced cells exhibited a 1.8- and 1.9-fold increases in H3-DPCs, respectively, compared with FA-treated WT cells (Figures 3G and 3H).

### *Sprtn* silencing increases *tdp1* expression in zebrafish embryos and human cell

To investigate the interplay between TDP1 and SPRTN at the gene expression level, we quantified their mRNA levels under different conditions of gene silencing and DPC induction. In zebrafish embryos, knockdown of *sprtn* resulted in a strong 2.2-fold increase in *tdp1* expression (Figure 4D), whereas this induction was much weaker in RPE1 cells where *SPRTN* silencing increased *TDP1* expression by 1.2-fold (Figure 4A).

**Figure 4.**
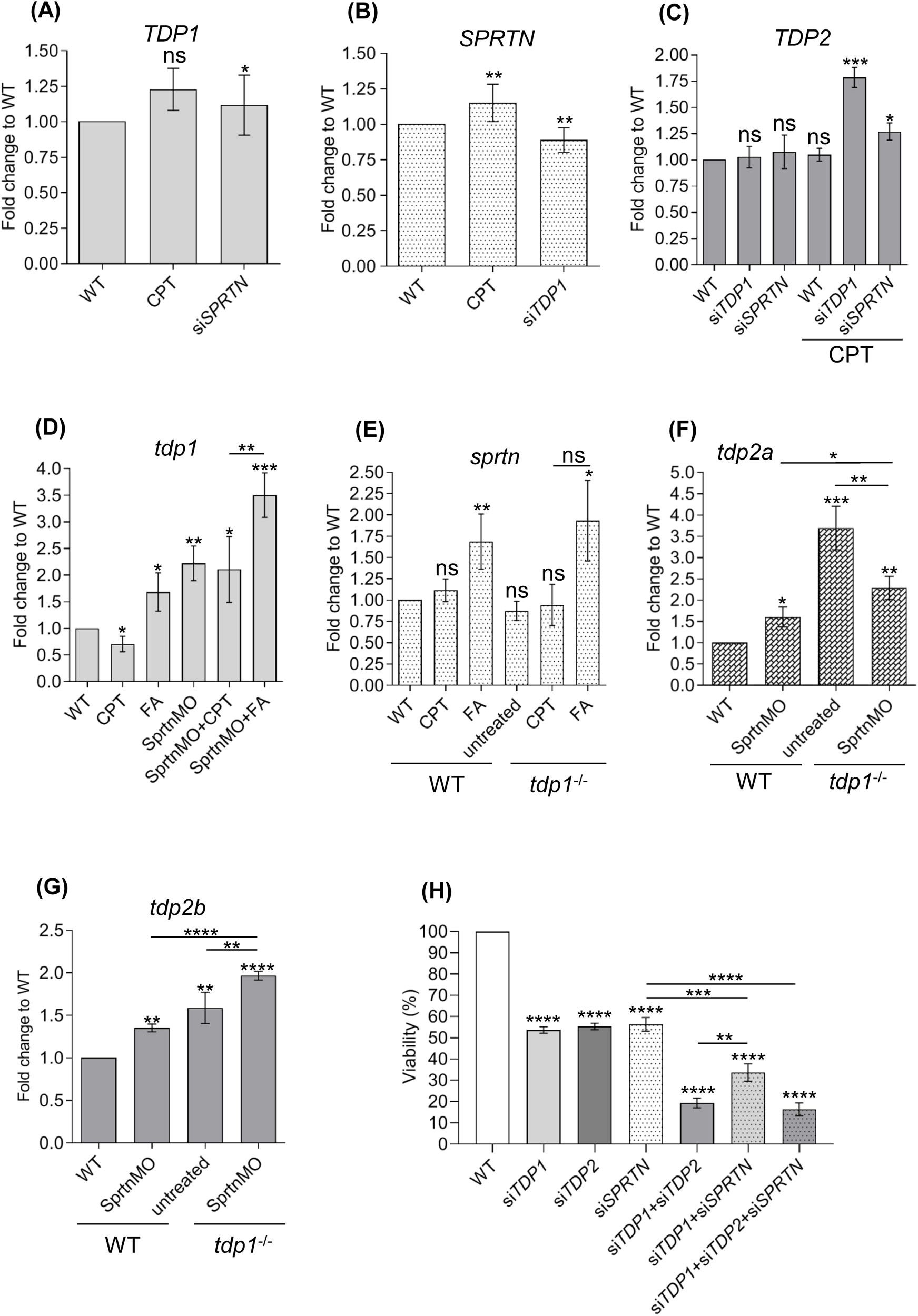
Effects of *TDP1* and *SPRTN* deficiency on *TDP1*, *SPRTN*, and *TDP2* mRNA expression levels in RPE1 cells and zebrafish embryos and decrease of cell viability. A) Expression levels of *TDP1* in RPE1 cells after *SPRTN* silencing and CPT exposure (50 nM, 1 hour) B) *SPRTN* levels decrease after *TDP1* silencing in CPT-treated RPE1 cells. C) *TDP2* significantly increases after *TDP1* silencing in CPT-treated RPE1 cells. D) Zebrafish *tdp1* expression levels significantly increase in embryos after *sprtn* knockdown. D) *Sprtn* expression in WT and *tdp1* mutant embryos before and after CPT (10 µM, 1h) and FA (1 mM, 20 min) treatment. F) *Tdp2a* expression is significantly increased in *tdp1* mutants before and after *sprtn* silencing and in WTs after *sprtn* silencing. (G) *Tdp2b* expression significantly increases in *tdp1* mutants before and after *sprtn* silencing and in WTs after *sprtn* silencing. Results are presented as fold changes to WT (mean ± SD) from four biological replicates. H) MTT viability assay after *TDP1*, *SPRTN* and *TDP2* gene silencing. All measurements were normalized to WT from three different experiments. Corresponding silencing efficiencies are shown in supplementary Figure S3 (mean ± SD; n = 3 independent experiments). Unpaired t-tests were performed with GraphPad Prism, with significant shown as * (*p* < 0.05), **(*p* < 0.01), ***(*p* < 0.001), or ****(*p* < 0.0001).

Brief acute exposure of embryos to CPT (10 µM, 1h) which strongly induced Top1-DPCs (2.8-fold) (Figures 2A and 2B) resulted in a 29% (0.7-fold) decrease in *tdp1* expression (Figure 4D). In contrast, a lower dose of CPT (1 h, 50 nM) in RPE1 cells, which induced TOP1-DPCs by 2-fold (Figure 2E and 2F), did not significantly alter *TDP1* expression (Figure 4A). The combination of *sprtn* silencing and CPT treatment had no further effect on *tdp1* expression compared to *sprtn*-silenced non-treated embryos (Figure 4D). In contrast to the effects caused by CPT, treatment of WT embryos with FA (30 min, 5 mM) increased *tdp1* expression by 1.7-fold (Figure 4D). This effect was even more pronounced in FA-treated *sprtn*-silenced embryos, in which a 3.5-fold increase in *tdp1* expression was observed (Figure 4D).

The expression levels of *sprtn* in embryos were similar in WT and mutant embryos and did not change significantly after CPT exposure. However, FA increased *sprtn* expression 1.5-fold and 1.8-fold in WT and *tdp1* mutant embryos, respectively (Figure 4E). In RPE1 cells, *TDP1* silencing caused a 10% decrease in *SPRTN* expression (Figure 4B), whereas CPT exposure increased *SPRTN* expression by 1.2-fold (Figure 4B). Silencing and knockdown efficiencies are shown in Figures S2A, S2B and S2C.

### *Tdp2* expression increases in TDP1-deficient RPE1 cells and zebrafish embryos

In the absence of TDP1, TDP2 can participate in the Top1-mediated DNA damage in cultured DT40 cells and *in vitro* (Ledesma et al., 2009; Zeng et al., 2012). Therefore, we wanted to determine whether *TDP2* expression increases when TDP1 is depleted in cells and embryos. In RPE1 cells, *TDP2* expression remains the same after silencing of *TDP1* and *SPRTN* without exposure to DPC inducers. However, when cells are treated with CPT (50 nM, 1 h), a TOP1-DPC inducer, *TDP2* expression increases strongly (1.8-fold) in *TDP1*-silenced cells and moderately (1.3-fold) in *SPRTN*-silenced cells (Figure 4C). These data suggest that TDP2 may help overcome camptothecin-induced DNA damage in the absence of TDP1 in human cells. Because zebrafish has two *tdp2* orthologs, *tdp2a* (gene ID: 101887157) and *tdp2b* (gene ID: 553516), we performed qPCR analysis for both genes. In the absence of Tdp1, the expression of both genes increased significantly: *tdp2a* by 3.7-fold and *tdp2b* by 1.6-fold (Figures 4F and 4G). Expression of *tdp2a* also increased after *sprtn* knockdown, by 1.6-fold in WT and 2.3-fold in *tdp1* mutants. The effect of *sprtn* silencing on *tdp2b* expression is similar to the pattern observed for *tdp2a*, with a 1.4-fold increase in WT and a 2.1-fold increase in *tdp1* mutants (Figure 4G). In contrast, the expression pattern of the two *tdp2* orthologs in *tdp1* mutants is different: silencing of *sprtn* in mutants strongly decreased *tdp2a* expression compared with nonsilenced mutants (Figure 4F), whereas the pattern is reverse with respect to *tdp2b* expression, where silencing of *sprtn* causes an increase in *tdp2b* mRNA levels (Figure 4G).

### TDP1 and SPRTN deficiency affects cell viability

We observed a significant reduction in cell density after silencing of *TDP1* and *SPRTN* in RPE1 cells, so we quantified this effect using the MTT assay. Considering that TDP2 could compensate for the loss of TDP1 (Ledesma et al., 2009), we also investigated the effects of TDP2 deficiency in combination with the lack of TDP1 and SPRTN on cell survival. Individual silencing of *TDP1*, *TDP2*, or *SPRTN* decreased cell viability by 50% (Figure 4H). Simultaneous silencing of *TDP1* and *SPRTN* further decreased cell viability by 68% (p < 0.0001). Interestingly, the effect was most pronounced when both *TDP1* and *TDP2* were silenced, and viability decreased by 80%. We observed a similar effect, an 84% decrease in viability, after all three genes were silenced (Figure 4H). The experiment was performed with three independent biological replicates, and silencing efficiencies were measured for each condition (Figure S3).

### Loss of Tdp1 leads to DPC accumulation in zebrafish embryos

*Tdp1* mutants have significantly higher endogenous DPC levels than WT embryos. The change is 1.4-fold increase and is statistically significant (Figures 5A and 5B, n = 4). All experiments in embryos were repeated 4-6 times (biological replicates), because the results showed higher variability compared to experiments in RPE1 cells. CPT treatment had no effect on total DPC levels in WT embryos, but caused a significant increase (1.7-fold) in *tdp1* mutants (Figures 5A and 5B). In contrast, the general DPC inducer, FA, caused a similar increase in total DPCs independent of Tdp1 deficiency: 2.2-fold in WTs and 2.1-fold in *tdp1* mutants (Figures 5A and 5B).

**Figure 5.**
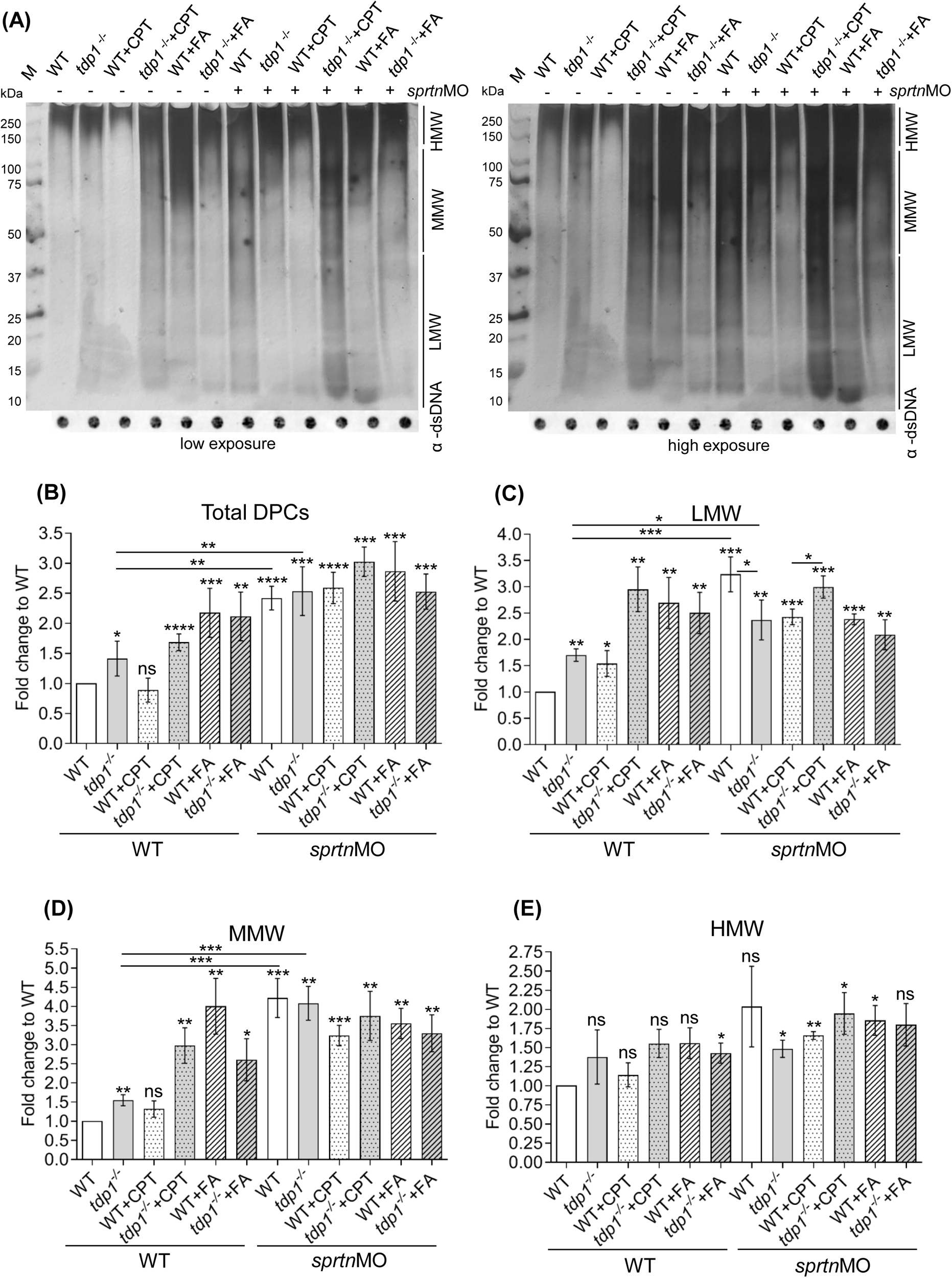
DPC analysis in Tdp1 and Sprtn deficient embryos under physiological conditions and after CPT (10 µM, 1h) and FA (5 mM, 30 min) treatment. A) DPCs were isolated from 2 dpf embryos using the RADAR assay (30 embryos per condition, n = 4), resolved on the SDS acrylamide gel, and stained with silver (left panel - low exposure, right panel - high exposure). Dot-blots showing DNA loading controls for DPC analysis prior to benzonase treatment are shown below (DPC equivalent of 200 ng of total DNA was loaded per well). B) Quantification of A. Quantifications of LMW DPCs (protein size < 40 kDa) (C), MMW DPCs (40 kDa to 150 kDa) (D) and HMW (>150 kDa) (E) from (A). Data represent mean fold change to WT ± SD (n = 4), statistical significance was established using an unpaired Student’s t-test (* (*p* < 0.05), ** (*p* < 0.01), *** (*p* < 0.001), and **** (*p* < 0.0001)).

Following the analysis of the endogenous and exogenous total DPC levels in *tdp1* mutants, we investigated the interplay of Tdp1 and Sprtn in DPC removal at the *in vivo* level. Knockdown of *sprtn* resulted in a strong and significant increase in total DPC levels in both, WT and *tdp1* mutant embryos (2.4-fold and 2.5-fold, respectively) (Figures 5A and 5B). Compared with untreated WT embryos, CPT exposure increased total DPCs in *sprtn*-silenced embryos 2.6-fold in WT and 3.1-fold in *tdp1* mutants (Figure 5B). Also, *sprtn* silencing caused a significant additive effect on total DPC increase (p < 0.001) in CPT-treated WT and mutant embryos (Figure 5B). In contrast, the effect of *sprtn* silencing was not significant in either WT (p > 0.05) or mutant embryos (p > 0.05) when exposed to the general DPC inducer, FA (Figure 5B).

Analysis of total cellular DPCs revealed important details about the functions of TDP1 and SPRTN in DPC removal and the effects of DPC inducers. However, to determine which DPCs are most affected by these changes, DPCs were divided into three subgroups: High Molecular Weight (HMW > 151 kDa), Medium Molecular Weight (MMW, 40 kDa to 150 kDa), and Low Molecular Weight (5 - 40 kDa). We are aware that this categorization is not ideal, but it can provide valuable additional information for the analysis of cellular DPCs compared with the analysis of total DPCs. More detailed analysis revealed that the 1.4-fold increase in total endogenous cellular DPCs in the *tdp1* mutants was indeed due to the increase in low and medium molecular weight DPCs, whereas high molecular weight DPCs were not affected (Figure 5A, high exposure). Specifically, Tdp1-deficient embryos accumulated 1.7-fold more endogenous LMW DPCs and 1.6-fold more endogenous MMW DPCs than WT embryos (Figures 5C and 5D). CPT treatment further increased LMW DPC levels: 3.1-fold in mutants and 1.5-fold in WTs (Figure 5C). A different pattern was observed after induction of general DPCs by FA, where levels of LMW DPCs increased 2.7- to 2.5-fold in both WT and mutant embryos (Figures 5A and 5C). As expected, treatment with FA had strong effects on LMW DPC levels considering that most of cellular DPCs are histones (Kiianitsa & Maizels, 2020). Unexpectedly, LMW DPCs were also induced by CPT treatment (1.5-fold), although not as strongly as after FA treatment (Figures 5A and 5C). When *sprtn* was knocked down, WT embryos accumulated more LMW DPC (3.1-fold) than mutants (2.4-fold) (Figures 5A and 5C). CPT treatment further increased LMW in mutant embryos 3.1-fold, but had no effect on LMW levels in WT embryos after knockdown of *sprtn*. LMW levels were not further affected by FA treatment when *sprtn* was silenced in WT or *tdp1*-deficient embryos (Figure 5C).

CPT treatment of *tdp1* mutants strongly increased the levels of MMW DPCs (by 2.9-fold), in contrast to a slight statistically nonsignificant change in WT embryos (Figures 5A and 5D). FA increased MMW DPCs by 2.6-fold in mutants and by 4.1-fold in WT embryos, showing a similar pattern of induction as LMW but with much stronger absolute changes. MMW DPCs increased 4.1-fold in both WT and *tdp1* mutants after *sprtn* knockdown (Figures 5A and 5D). MMW DPCs in WT embryos and *tdp1* mutants were not additionally affected by CPT or FA treatment in *sprtn* knockdowns (Figures 5A and 5D).

Tdp1 deficiency, *sprtn* knockdown, and exposure to CPT or FA had the least effect on HMW DPCs. However, some of the effects were still pronounced. HMW DPCs increased following *sprtn* knockdown by 1.5-fold in *tdp1* mutant embryos (*p* < 0.05) and by 2-fold in WT embryos (p > 0.05) (Figures 5A and 5E). HMW DPCs increased 1.9-fold in the *tdp1* mutant and 1.6-fold in WT when *sprtn* knockdown was combined with CPT treatment. Independent of Tdp1 deficiency, knockdown of *sprtn* showed a similar induction by 1.8-fold in both the *tdp1* mutant and WT embryos which were treated with FA (Figures 5A and 5E). Minor variations in HMW DPCs observed between the *tdp1* mutant and WT embryos with and without CPT or FA treatment were not statistically significant.

### *TDP1* silencing causes DPC accumulation in human cells

DPC levels were quantified in RPE1 cells after *TDP1* and *SPRTN* silencing. Silencing of *TDP1* alone caused a small, but statistically significant increase in total DPCs (1.2-fold) (Figures 6A and 6B), whereas silencing of *SPRTN* caused a bigger, 1.6-fold increase in DPC levels (Figures 6A and 6B, n = 4). When both, *TDP1* and *SPRTN* were silenced (Figure S2C), we observed an additive effect: a 2-fold increase in total cellular DPCs (Figures 6A and 6B). Considering that SPRTN is involved in the removal of very diverse DPCs, ranging from low molecular weight (LMW) proteins such as histones to bulky high molecular weight (HMW) proteins such as topoisomerases (Vaz et al., 2016), we further investigated the size distribution of the isolated DPCs. *TDP1* silencing increased low and medium molecular weight (LMW and MMW) DPCs by 1.3- and 1.5-fold, respectively (Figures 6A right panel, 6C and S4A). The effect of the increase is not as strong as that of *SPRTN* silencing, which showed an increase of 2.6-fold in the LMW region and 1.9-fold in the MMW region (Figure 6C). The silencing combination showed an additive effect on the increase in LMW and MMW DPCs (3.2- and 2.2-fold, respectively). The silencing combination also showed a 1.7-fold increase in HMW DPCs, in contrast to single gene silencing, where no increase was observed (Figures 6A and 6C).

**Figure 6.**
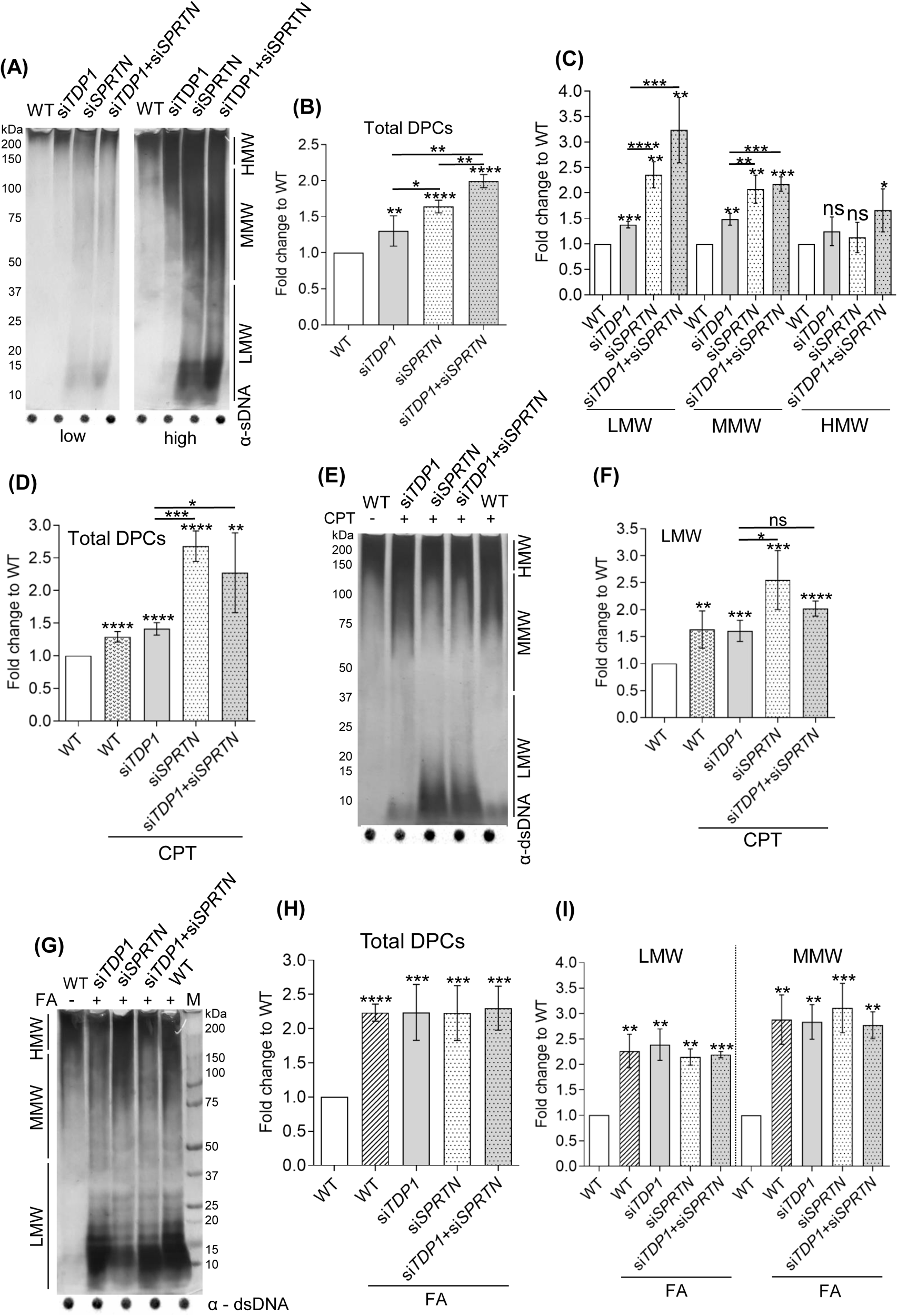
DPC analysis in RPE1 cells after *TDP1* and *SPRTN* gene silencing and after CPT (50 nM, 1h) and FA (1 mM, 20 min) treatment. Silencing was carried out for 72 hours prior to collection, and the efficiency of each condition was confirmed using qPCR (Figures S2C). A) DPC isolates from untreated cells resolved on the SDS acrylamide gel, and stained with silver (left panel - low exposure, right panel - high exposure). Dot-blots showing DNA loading controls are shown below. B) Quantification of A. C) Quantification of LMW, MMW and HMW DPCs from (A) normalized to non-treated WT cells from four independent experiments (n = 4) D) Quantification of E (n = 3). E) DPC isolates from CPT-treated cells resolved on the SDS acrylamide gel and stained with corresponding DNA loading controls shown below. F) LMW DPCs (quantification from D). G) DPC isolates from FA-treated cells resolved on the SDS acrylamide gel and stained with silver with corresponding DNA loading controls. H) Quantification of G (n = 3). I) LMW and MMW DPC levels quantified from (G), a DPC equivalent of 200 ng total DNA was loaded per condition. All conditions were normalized to WT and statistical analysis was performed with GraphPad Prism software using an unpaired t-test (* (*p* < 0.05), ** (*p* < 0.01), *** (*p* < 0.001), or **** (*p* < 0.0001)).

After analyzing DPC accumulation in untreated cells, we quantified DPCs after exposure to camptothecin (CPT), a specific TOP1-DPC inducer, and after exposure to formaldehyde (FA), a strong general DPC inducer. We used 50 nM CPT (1 h, 37 ᵒC) in serum-free medium which induces TOP1-DPCs without the occurrence of double-strand breaks (Ray Chaudhuri et al., 2012) and 1 mM FA in ice cold serum-free medium (20 min, 37 ᵒC) which induces DPCs and probably also single- and double-strand breaks based on the data from HEK293 cells (Mórocz et al., 2017). Treatment with CPT had a similar effect on DPC accumulation as did *TDP1* silencing: total DPCs increased by 1.3-fold, with the largest effect on LMW DPCs with a 1.6-fold increase (Figures 6D, 6E and 6F). When CPT was added to *TDP1*-silenced cells, a 1.4-fold increase in total DPCs was observed (Figure 6D), again with the largest effect on LMW with a 1.6-fold increase (Figures 6E and 6F). CPT exposure of *SPRTN*-silenced cells caused very strong DPC accumulation (2.7-fold compared with untreated WT cells), again with the largest effect on LMW DPCs of 2.-fold increase (Figures 6D, 6E and 6F). In contrast to untreated cells, treatment with CPT after double silencing had no additional effect on DPC accumulation (Figures 6D and 6E). Similar to LMW DPCs, treatment with CPT resulted in a slight 1.3-fold increase in MMW DPCs (Figure S4B). Interestingly, MMW DPCs in cells with silenced *TPD1*, *SPRTN*, or *TDP1* and *SPRTN* were equally affected whether CPT was applied or not (Figure S4B). The CPT exposure had the least effect on HMW DPCs (Figure S4B).

In contrast to the DPC response to CPT treatment, the pattern of DPC accumulation after FA treatment was very different. FA treatment increased total DPC levels by 2-fold in all samples regardless of which gene was silenced (Figures 6G and 6H). In all samples, FA treatment had the greatest impact on LMW and MMW DPCs, which increased by 2.3 and 2.8-folds on average in comparison to untreated WT cells (Figure 6I). Proteins of high molecular weight were least affected by FA treatment and showed no statistically significant difference in comparison to WT (Figure S4C).

## Discussion

We show that TDP1 is a key factor for the repair of histone H3- and TOP1-DPCs, while SPRTN is crucial for the repair of multiple cellular DPCs including TOP1- and H3-DPCs in human cells and in zebrafish (Figure 7). We further demonstrate resolution of H3-DNA crosslinks depends on upstream proteolysis by SPRTN and subsequent peptide removal by TDP1 in cells and embryos (Figure 7). In contrast to H3-DPC repair, where SPRTN and TDP1 work together, we show that they function in separate pathways in the repair of endogenous TOP1-DPCs (Figure 7). However, after exposure of human cells to clinically relevant concentrations of camptothecin, SPRTN and TDP1 act epistatically in the resolution of total DPCs, histone H3- and TOP1-DPCs (Figure 7).

**Figure 7.**
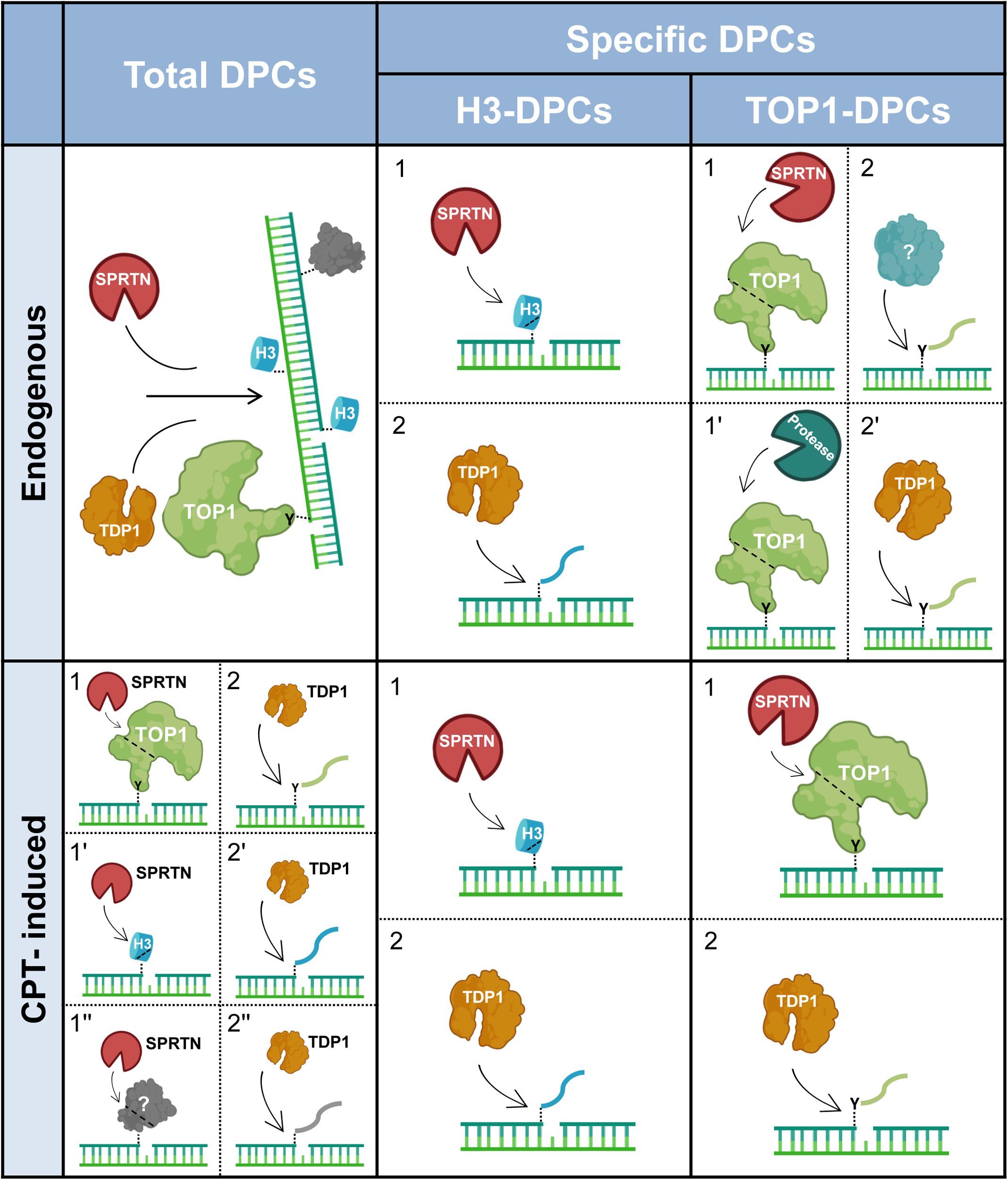
Model of coordinated action of SPRTN and TDP1 in DNA-protein crosslink repair in human cells and zebrafish model. SPRTN is a general DPC protease that cleaves a wide spectrum of crosslinked proteins, whereas TDP1 removes protein residues bound to the 3’ end of the ssDNA break. Under physiological conditions, these two proteins function independently to resolve total DPCs, including specific TOP1-DPCs. Importantly, resolution of endogenous histone-DPCs originating at abasic (AP) sites depends on SPRTN-mediated proteolysis followed by TDP1 phosphodiesterase activity, by which the crosslinked peptide residue is removed from the DNA backbone. In response to the DNA-damaging agent camptothecin, an epistatic relationship between SPRTN and TDP1 is required for the successful removal of histones and TOP1-DPCs as well as other DPCs at the 3’ ends of ssDNA breaks. The model was created with BioRender.com.

To study DPC repair *in vivo*, we established the toolbox for using the zebrafish animal model in DPC research, which includes optimization of protocols for DPC isolation and detection from embryonic tissue, generation of a Tdp1-deficient zebrafish strain, optimization of morpholino-mediated *sprtn* knockdown, and development of Tdp1 antibody which recognizes zebrafish ortholog. Understanding how DPC repair factors function in organisms is only possible through research in animal models, and we hope that the Tdp1-deficient zebrafish strain we have created will be a valuable resource for future studies of DPCR in specific tissues.

We analysed the degree of conservation of TDP1 between humans, mice and zebrafish and found that zebrafish is an acceptable vertebrate model for studying TDP1 function. The high degree of evolutionary conservation of one-to-one orthology across all domains of life (Figures S1A and S1B) and the very high structural similarity between zebrafish and human orthologs (Figure 1A) confirm the importance of TDP1 for DNA repair throughout evolution.

High mRNA expression levels of *tdp1* and *sprtn* during embryonic development indicate that both repair factors are important in the developing embryo (Figure 1F). This result is not surprising given rapid rate of cell division and transcription during this period and thus, the high demand for precise DNA repair (Keller, 2013). *TDP1* is highly expressed in human testis (Uhlén et al., 2015) and zebrafish and mouse gonads (Figures 1G and 1H), suggesting its protective role in germ cells. A similar mRNA tissue expression pattern of *Tdp1* in zebrafish and laboratory mouse (Figures 1G and 1H) suggests that zebrafish would be a good model for future investigation of the tissue-specific function of TDP1 in DPCR. Comparison with human expression data is currently not possible because the mRNA expression dataset available in the Human protein Atlas is heavily biased toward analysis of older individuals (Uhlén et al., 2015).

With respect to the canonical TDP1 substrate, TOP1, we found that TDP1 is crucial for the removal of TOP1-DPCs, both at the organism level and in RPE1 cells (Figure 2). Consistent with previous findings in HEK293 human cells (Fielden et al., 2020), the silencing of *TDP1* in RPE1 cells resulted in increased TOP1-DPC levels (Figures 2E and 2F). In contrast, knock-out of *TDP1* in RPE1 did not lead to TOP1-DPC increase (Meroni et al., 2022), suggesting adaptive mechanisms in permanent knock-out as opposed to temporary gene silencing. Given the importance of TOP1 inhibition in cancer therapy (Fengzhi Li, 2017; Martino et al., 2017), the role of TDP1 in TOP1-DPC repair has been extensively investigated *in vitro* and in cell cultures (Gao et al., 2014; Heidrun & James, 2011; Pouliot et al., 1999; Yang et al., 1996), but data on the vertebrate models is sparse (Hirano et al., 2007; Katyal et al., 2007; Zaksauskaite et al., 2021). Using zebrafish model, we show that Tdp1 deficiency leads to a significant 4.2-fold increase in endogenous Top1-DPCs (Figures 2A and 2B), demonstrating that TDP1 is crucial repair factor for the resolution of TOP1-DPCs. Previous study by Zaksauskaite *et al*. (2021) did not report difference in Top1-DPC levels between wild-type and Tdp1-deficient zebrafish embryos. The discrepancy is likely due to the different approaches to DPC isolation, as the RADAR assay used in this study is more specific and sensitive than the CsCl fractionation method used in the previous study (Kiianitsa & Maizels, 2013, 2014, 2020). Expression analysis of *tdp2* showed a significant increase in *tdp2* when *tdp1* was absent in embryos suggesting that TDP2 may compensate for loss of TDP1 *in vivo*, supporting previous observations from cell culture and *in vitro* experiments (Ledesma et al., 2009; Zeng et al., 2012). Detailed insights into TOP1-DPC repair at the organism level are important for biomedicine, because TOP1-DPC inducers, the camptothecin derivatives irinotecan and topotecan, are used to treat various cancers including ovarian and colon cancers, small-cell lung cancer, central nervous system tumors, and sarcomas (Martino et al., 2017). Combination therapies with TOP1 and TDP1 inhibitors could significantly improve current clinical protocols, and thus it is essential to know how TDP1 functions in the repair of DPCs at the organism level.

It is also very important to understand how histone-DPCs are repaired because they are very abundant under physiological conditions. More than 10,000 abasic sites are generated daily (Lindahl, 1993), and about 10% of these lead to the formation of DPCs, most of which are histone-DPCs (Kiianitsa & Maizels, 2020). This suggests that hundreds, possibly even thousands of histones are cross-linked to abasic sites in each individual cell every day (Ren et al., 2019). Induction of histone-DPCs may be a promising new avenue to explore in the treatment of cancer. Our discovery that TDP1 is a critical factor in histone-DPC repair (Figure 3) further highlights TDP1 as an important drug target. Our results support recent observations of Wei *et al*. (2022), who showed that TDP1 can remove histone H2B and H4 from AP sites *in vitro*, and provide evidence for a novel, TDP1-dependent repair pathway for histone-DPC resolution *in vivo*. We also observed that loss of Tdp1 in zebrafish embryos and RPE1 cells results in a significant increase in cellular DPCs that cannot be attributed solely to the increase in H3- and TOP1-DPCs (Figures 5 and 6), suggesting that TDP1 has multiple DPC substrates. On the other hand, the increase in low molecular weight DPCs in *tdp1* mutant embryos and TDP1-deficient RPE1 cells (Figures 5C and 6C) is most likely due to the increase in histone-DPCs, considering that endogenous H3-DPCs accumulate strongly in Tdp1-deficient embryos and human cells (Figure 3). The function of TDP1 in the repair of cellular DPCs and histone-DPCs has not been previously investigated and our results should be considered in the development of TDP1 inhibitors for cancer therapy (Sun et al., 2020).

Regarding the role of SPRTN protease in DPCR, we show that SPRTN is a crucial protease for the resolution of multiple cellular DPCs *in vivo*, and in particular for the removal of low- and medium-molecular weight DPCs (Figures 5 and 6). By analysing DPCs in SPRTN-deficient embryos, we provide the first evidence of how DPC levels are affected in an organism. Compared with embryos, the effect of SPRTN deficiency was somewhat weaker in RPE1 cells. We have shown that SPRTN is critical for TOP1-DPC repair at the organism level (Figure 2C and 2D), supporting previous studies in cell culture that showed an increase in TOP1-DPC levels after SPRTN silencing (Maskey et al., 2017; Stingele et al., 2014; Vaz et al., 2016) and that SPRTN proteolyzes TOP1 *in vitro* (Vaz et al., 2016). Our results also show that SPRTN plays an important role in resolving H3-DPCs *in vivo*, as knockdown of *sprtn* in zebrafish resulted in a 5.7-fold increase in H3-DPC levels (Figure 3A and 3B), supporting *in vitro* data characterising H3 as a substrate of SPRTN (Vaz et al., 2016). Therefore, our study finally demonstrates the crucial role of SPRTN in resolving DPCs at the organism level and highlights SPRTN as a promising chemotherapeutic target.

We characterized the interplay of TDP1 and SPRTN in DPC resolution under physiological conditions. SPRTN and TDP1 are known to play distinct roles in DPC repair. SPRTN is required for initiating repair of many crosslinked proteins (Fielden et al., 2018), whereas TDP1 has been specifically associated with repair of TOP1-DPCs (Kawale & Povirk, 2018). Indeed, our results in RPE1 cells suggest a non-epistatic relationship between these two proteins in the repair of total cellular DPCs (Figures 6A, 6B and 7) and in the repair of endogenous TOP1-DPCs (Figures 2E, 2F and 7). We observed a similar pattern in embryos, but the changes were not statistically significant in this case because of higher variability between experiments (Figure 2C and 2D). Since it is known that TDP1 cannot process TOP1-DPCs alone (Heidrun & James, 2011), we hypothesize that another protease is involved in the upstream proteolysis of endogenous TOP1-DPCs (Figure 7). Possible candidates include DDI1, DDI2, FAM111A, and the proteasome (Dirac-Svejstrup et al., 2020; Kojima et al., 2020; Larsen et al., 2019; Serbyn et al., 2020). Unlike in endogenous TOP1-DPC repair, we show that SPRTN and TDP1 work together in the resolution of endogenous H3-DPCs in zebrafish and in human cells (Figures 3A, 3B, 3E and 3F), suggesting that SPRTN is the main protease acting upstream of TDP1-mediated peptide removal in the resolution of histone-DPCs at AP sites (Figure 7). Our results are the first to show that SPRTN proteolysis is required for histone-DPC resolution *in vivo*. It is worth noting that *in vitro*, TDP1 can remove H2B and H4 crosslinks without the requirement of upstream proteolysis (Wei et al., 2022). Known discrepancies between in vitro and in vivo data further emphasizes the urgent need for the experimental data from the animal models. The interplay between SPRTN and TDP1 is also evident at the level of mRNA expression, where silencing of *SPRTN* increases *TDP1* expression in embryos and cells (Figure 4A and 4D). Both proteins are critical for normal cell function, as RPE1 cells with silenced *TDP1* and *SPRTN* exhibit a 68% reduction in viability (Figure 4H). We suggest that this phenotype is the result of impaired DPCR resulting from the absence of TDP1 and SPRTN (Figures 2, 3, 5 and 6). It is known that *SPRTN*-silenced human cells exit S phase with abnormal replication intermediates (Mórocz et al., 2017) and *TDP1*-silenced cells exhibit an increase in ssDNA and dsDNA breaks (Fielden et al., 2020).

In contrast to physiological conditions, after exposure to CPT, when TOP1-DPCs are induced, we show that SPRTN and TDP1 act epistatically in resolving total DPCs in RPE1 cells (Figures 6A, 6B, 6D, 6E and 7). Our results are consistent with data from cell survival assays showing that simultaneous depletion of TDP1 and SPRTN in yeast (Stingele *et al*., 2014; Lopez-Mosqueda *et al*., 2016) and HeLa cells (Vaz et al., 2016) to a similar extent as depletion of either component alone leads to hypersensitivity to treatment with CPT, suggesting that SPRTN and TDP1 function in the same pathway for the repair of CPT-induced TOP1-DPCs. It is important to note that DPCR factors may behave differently under physiological conditions and under stress, when DPC load exceeds certain thresholds. In summary, our results suggest that repair of CPT-induced TOP1-DPCs relies on the SPRTN-TDP1 axis, in contrast to repair of endogenous TOP1-DPCs, in which TDP1 and SPRTN function in separate pathways (Figure 7). It is important to note that CPT also induces H3-DPCs in human cells and in zebrafish embryos (Figures 3, S2E and S2F), demonstrating for the first time that CPT is not specific for TOP1-DPC induction, as previously thought (Ramawat & Mérillon, 2013). Whether this effect is direct or indirect remains to be determined in future studies.

Formaldehyde is a potent crosslinker of various cellular proteins ranging in size from 10 kDa to over 200 kDa (Stingele *et al*., 2014; Lopez-Mosqueda *et al*., 2016; Ruggiano *et al*., 2021). Our results show that the interplay of SPRTN and TDP1 in DPCR is altered when cells and embryos are exposed to high acute doses of FA, compared with physiological conditions in which only endogenous DPCs are present. SPRTN and TDP1 act together (in epistasis) in the repair of FA-induced total cellular DPCs (Figures 5 and 6G, 6H and 6I) and in the repair of FA-induced H3- and TOP1-DPCs (Figures 2 and 3). Considering that large amounts of DPCs accumulate under these conditions, we hypothesize that SPRTN is fully activated and performs upstream proteolysis of many cross-linked proteins given its pleiotropic nature (Vaz et al., 2016), and that TDP1 is crucial for the resolution of H3- and TOP1-DPCs and other crosslinked protein residues at AP sites.

In summary, we have performed a comprehensive analysis of the role of TDP1 and SPRTN in the resolution of DPCs in zebrafish and human cells. Our results reveal the interplay of these repair factors in the resolution of cellular DPCs, H3- and TOP1-DPCs and introduce a novel TDP1-mediated repair pathway for histone-DPCs that highlights the epistatic relationship between upstream SPRTN proteolysis and downstream TDP1-mediated 3’ DNA end processing (Figure 7). Furthermore, we demonstrate the essential role of both proteins in the resolution of total cellular DPCs and TOP1-DPCs in human cells and in the animal model. Our results provide new insights into the complex DPC repair pathways and their implications for human disease and cancer treatments. Further research in this area will advance our understanding of DPC repair factors and their potential therapeutic applications. It is important to point out that mechanistic, *in vitro* studies are essential for understanding DPC repair processes, but that research in animal models is essential for understanding and contextualising the interplay of the various repair factors in the whole organism which is a prerequisite for translating the acquired knowledge for understanding and treating human diseases. The mechanism of action of TDP1 and SPRTN has been previously studied in detail, but how these repair factors function together in cells and tissues in DPCR was previously unknown.

## Acknowledgments

This work was supported by Croatian Science Foundation Installation Grant (UIP-2017-05-5258). M.P. research group is supported by Slovenian-Croatian Bilateral Research Project grant (IPS-2020-01-4225) and European Structural and Investment Funds STIM – REI project (KK.01.1.1.01.0003). Frozen mouse tissues were a kind gift from dr.sc. Tihomir Balog (Ruder Boskovic Institute, Croatia).

## Authors contribution

IA created and validated the transgenic zebrafish line, optimized DPC isolation protocols, performed synteny analysis and all experiments on cell lines and zebrafish embryos, CO optimized gene silencing in embryos and performed injection experiments, LV and LJ performed expression experiments, MP performed phylogenetic and structural analysis, designed and supervised the project. IA and MP wrote the manuscript.

## Supplementary Figure legends

**Figure S1.**
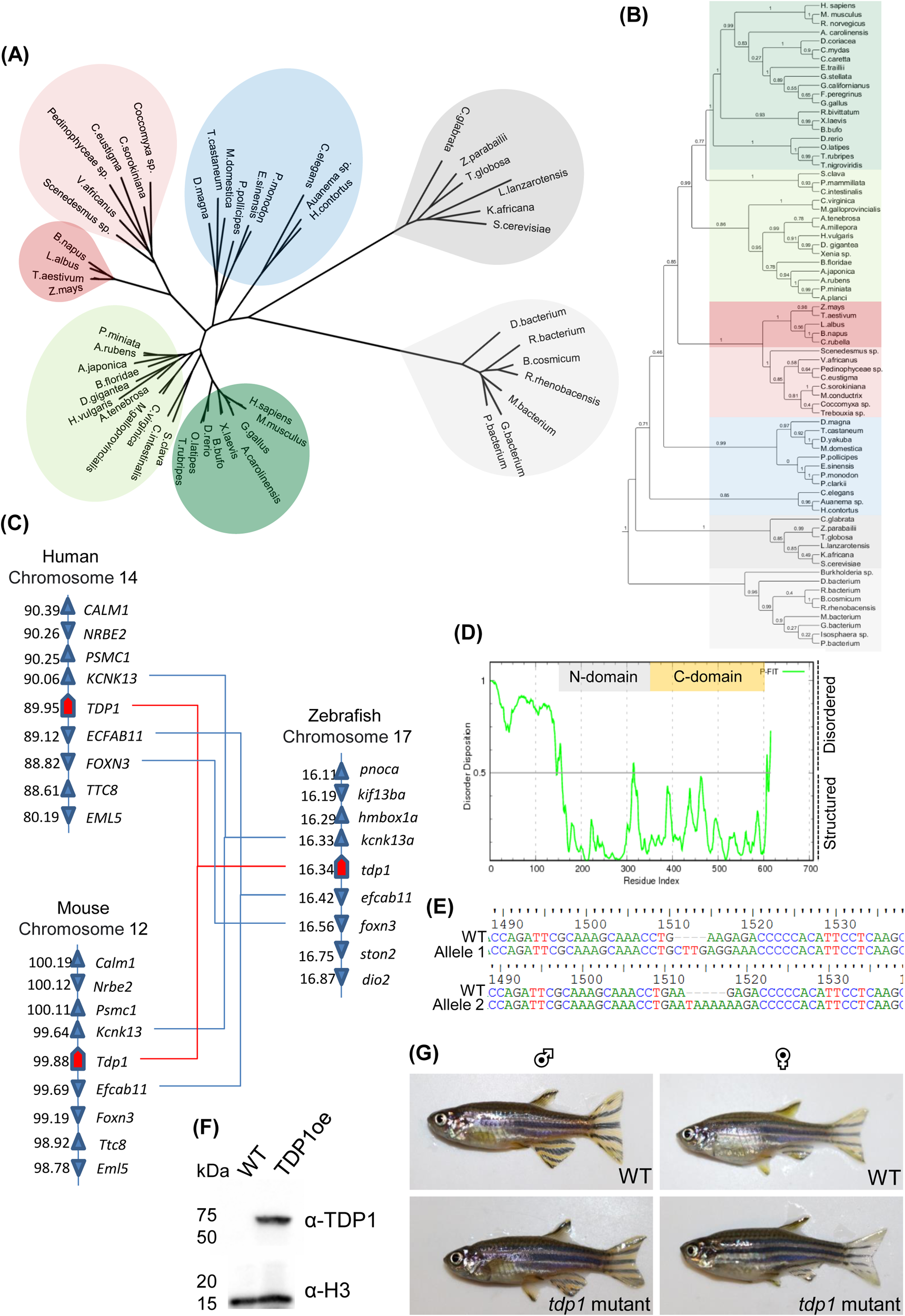
Phylogenetic, syntenic and sequence analysis of zebrafish Tdp1 and validation of *tdp1* mutant line. A) Phylogenetic tree of TDP1 orthologs. The vertebrate TDP1 cluster is shown in dark green, invertebrate cluster, which is phylogenetically closely related to vertebrate orthologs, is shown in light green, second invertebrate cluster is shown in blue. Plant and algae clusters are shown in dark and light red, respectively, while fungi and bacterial orthologs are shown in dark and light grey, respectively. B) Extended phylogenetic analysis of TDP1 proteins from bacteria to humans was performed using the Maximum Likelihood method with branch support Alrt values (Approximate likelihood-ratio test) are shown at tree nodes on a scale of 0-1, where 1 is maximum node confidence. C) Synteny analysis of zebrafish, human, and mouse *TDP1* performed using Genomicus (Louis et al., 2013). The numbers next to the gene names represent the megabase pair (Mbp) of each gene position on the chromosome. D) The plot of disorder disposition for zebrafish Tdp1 protein predicted by PONDR-FIT software. Values above 0.5 indicate likely disordered regions (N-terminal), while structured N- and C-domains are labelled in grey and yellow, respectively. F) Sequencing od *tdp1* mutant fish line that contains Tdp1 allele 1 bearing the insertion of CTTG after the position 1510.on gDNA and allele 2 bearing the TAAAA insertion after position 1512. on gDNA. F) Western blot showing that custom-made Tdp1 antibody recognizes the overexpressed human TDP1 recombinant protein in HEK293 cells. Histone H3 is shown as a loading control. G) WT and *tdp1* mutant adult fish (6 months old). Prior to imaging, adult fish were anesthetized with tricaine methanesulfonate (MS-222) (0.02% in aquarium water) for 10 minutes and left to recover afterwards. Images were captured using a Canon 250D DSLR camera.

**Figure S2.**
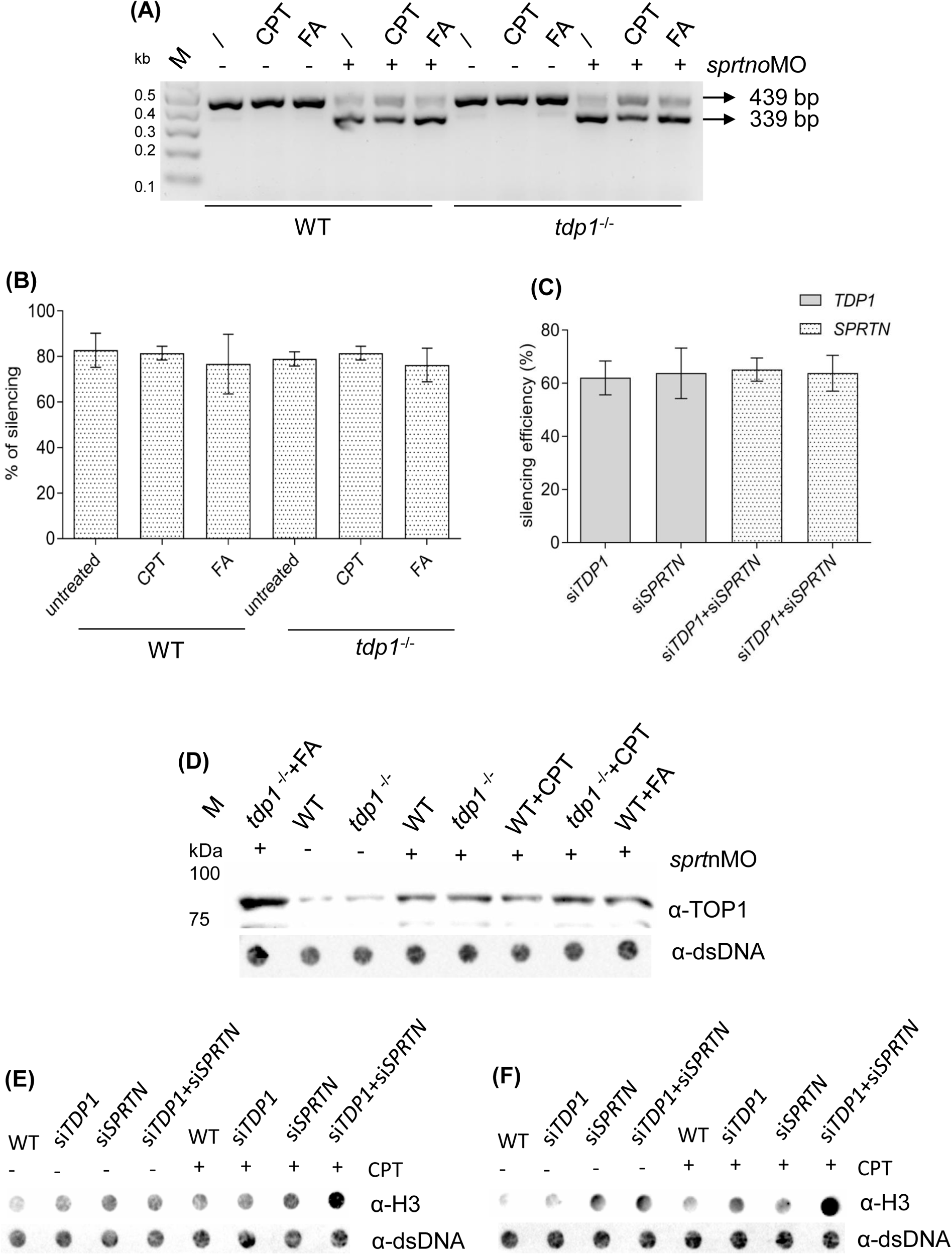
*Sprtn* and *TDP1* silencing efficiencies and biological replicates of Top1- and H3-DPC levels in zebrafish embryos and RPE1 cells. A) DNA gel electrophoresis showing the resolution of PCR reactions performed on cDNA isolated from two-day-old embryos after injection of exon-skipping morpholino into one cell stage embryos using primers listed in Table 4. B) Quantification of (A): *Sprtn* silencing efficiency was approximated as a reduction in WT bend, quantified by Image J and presented as a fold change to WT (mean ± SD) (n = 4). C) Efficiency of *TDP1* and *SPRTN* gene silencing in RPE1 cells. Cells were incubated with siRNAs listed in Table 1 for 72 hours. Silencing efficiency was quantified by qPCR: results are shown as the mean percent reduction in expression compared to untreated WT cells ± SD (n = 9). D) Western blot showing zebrafish Top1-DPCs in *tdp1* mutant embryos before and after silencing of *sprtn* and treatment with CPT (10 µM, 1 hour) and FA (5 mM, 30 min) used for quantification in Figure 2D. The equivalent of 1 µg DNA of total DPCs was loaded per sample. E) and D) Dot blots of H3-DPCs from two independent experiments after *TDP1* and *SPRTN* silencing before and after exposure to CPT (50 nM, 1h) with corresponding DNA loading controls used for quantifications shown in Figure 3F. The equivalent of 200 ng DNA of total DPCs was loaded per sample.

**Figure S3.**
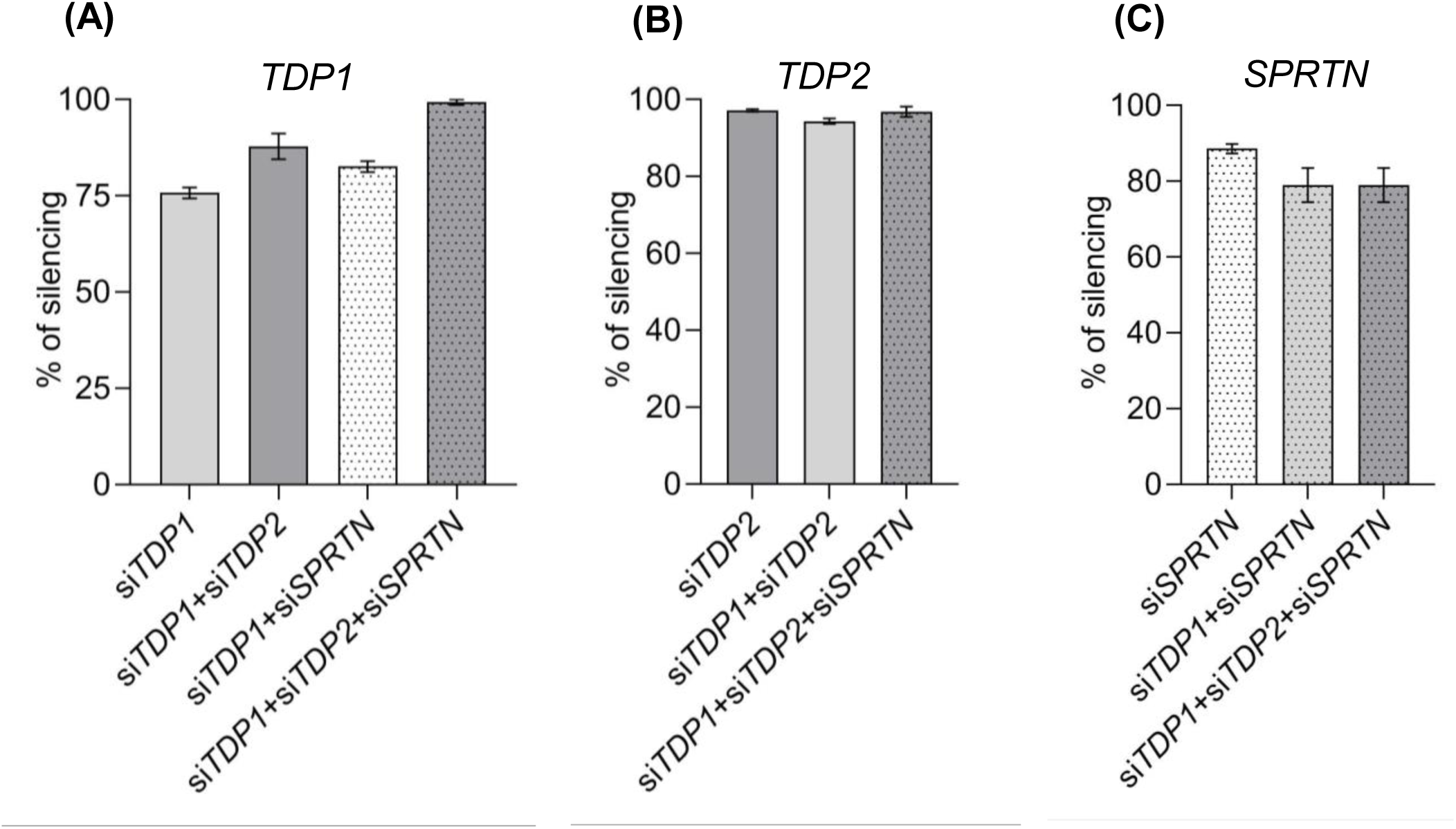
*TDP1*, *SPRTN*, and *TDP2* silencing efficiencies in the MTT cell viability assays. In parallel with the seeding of cells for the MTT viability assay shown in Figure 4H, cells were seeded for silencing verification using the same experimental conditions. Silencing efficiencies were determined using qPCR analysis and presented as the percentage reduction in expression to WT ± SD from three technical replicates.

**Figure S4.**
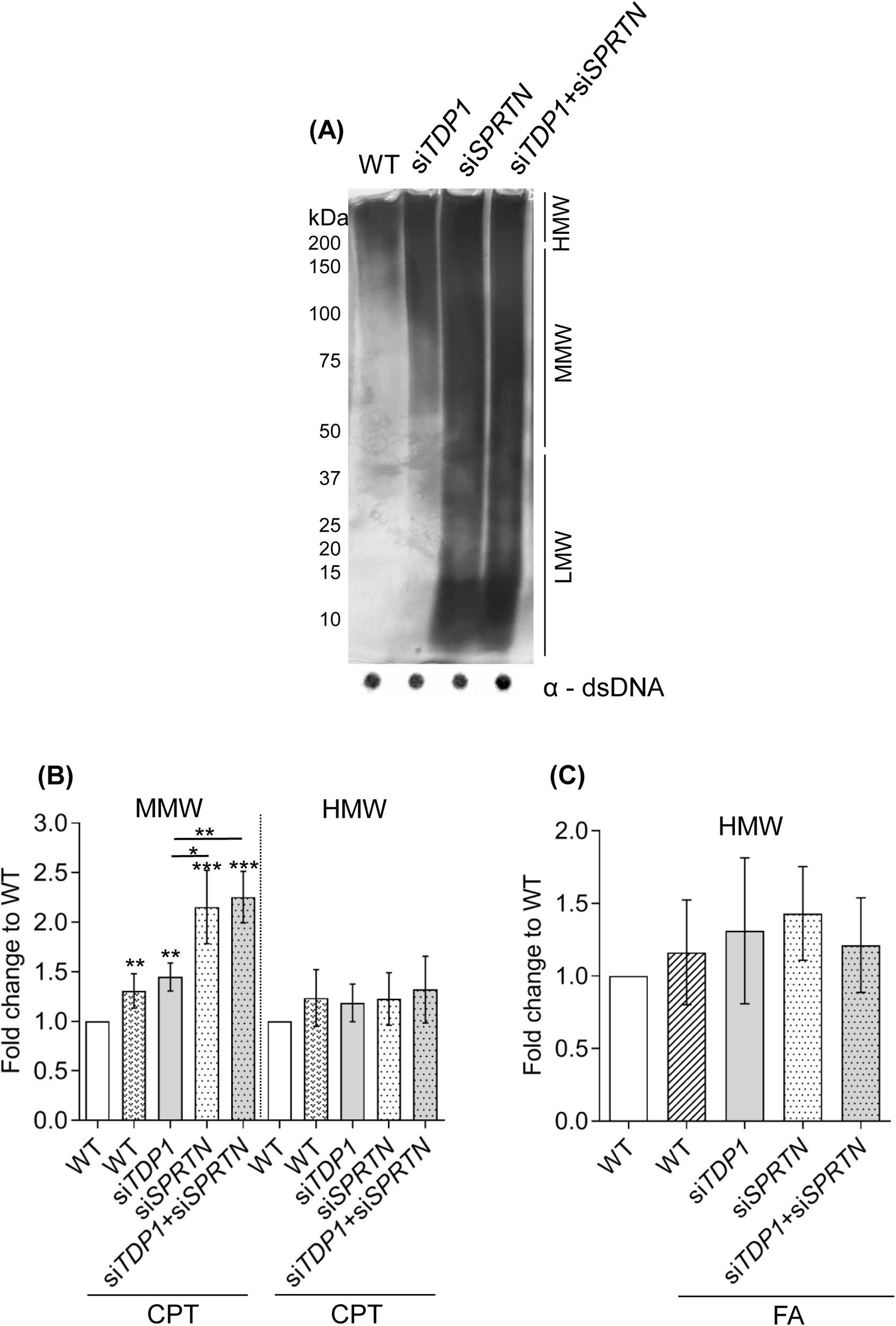
Total DPC analysis in RPE1 cells related to Figure 6. A) Higher exposure of silver stained gels shown in Figure 6A. B) Quantification of MMW and HMW DPCs from Figure 6E normalized to non-treated WT (mean ± SD; n = 3 independent experiments; * (*p* < 0.05), ** (*p* < 0.01), *** (*p* < 0.001), unpaired Student t-test). D) Quantification of HMW DPCs after exposure to FA shown in Figure 6G.

**Table S1.** Species names with corresponding protein accession numbers (NCBI database) used in phylogenetic analysis.

| Abbreviation | Accession no. | Species | Group |
| --- | --- | --- | --- |
| H.sapiens | NP_060789.2 | Homo sapiens | mammals |
| M.musculus | XP_030102367.1 | Mus musculus | mammals |
| R.norvegicus | XP_006240494.1 | Rattus norvegicus | mammals |
| G.gallus | XP_040528900.1 | Gallus gallus | birds |
| F.peregrinus | XP_005241969.1 | Falco peregrinus | birds |
| G.stellata | XP_009818907.1 | Gavia stellata | birds |
| G.californianus | XP_050754186.1 | Gymnogyps californianus | birds |
| E.traillii | XP_027758233.1 | Empidonax traillii | birds |
| A.carolinensis | XP_003214461.1 | Anolis carolinensis | reptiles |
| C.mydas | XP_037756745.1 | Chelonia mydas | reptiles |
| D.coriacea | XP_038262859.1 | Dermochelys coriacea | reptiles |
| C.caretta | XP_048709928.1 | Caretta caretta | reptiles |
| X.laevis | NP_001087094.1 | Xaneopus laevis | amphibians |
| R.bivittatum | XP_029453633.1 | Rhinatrema bivittatum | amphibians |
| B.bufo | XP_040268057.1 | Bufo bufo | amphibians |
| D.rerio | XP_700174.2 | Danio rerio | fish |
| O.latipes | XP_023809425.1 | Oryzias latipes | fish |
| T.rubripes | XP_003969465.3 | Takifugu rubripes | fish |
| T.nigroviridis | CAG03090.1 | Tetraodon nigroviridis | fish |
| B.floridiae | XP_035693489.1 | Branchiostoma floridiae | chordata |
| P.mammillata | CAB3266878.1 | Phallusia mammillata | tunicata |
| S.clava | XP_039264827.1 | Styela clava | tunicata |
| C.intestinalis | XP_002123899.2 | Ciona intestinalis | tunicata |
| P.miniata | XP_038045909.1 | Patiria miniata | echinodermata |
| A.japonica | XP_033125892.1 | Anneissia japonica | echinodermata |
| A.planci | XP_022107496.1 | Acanthaster planci | echinodermata |
| A.rubens | XP_033639145.1 | Asterias rubens | echinodermata |
| D.yakuba | XP_039227154.1 | Drosophila yakuba | insecta |
| M.domestica | XP_005187304.1 | Musa domestica | insecta |
| T.castaneum | XP_008191977.1 | Triboleum castaneum | insecta |
| D.magna | KZS04356.1 | Daphnia magna | crustacea |
| P.monodon | XP_037795115.1 | Penaeus monodon | crustacea |
| P.clarkii | XP_045608059.1 | Procambarus clarkii | crustacea |
| E.sinensis | XP_050707432.1 | Eriocheir sinensis | crustacea |
| P.pollicipes | XP_037088035.1 | Pollicipes pollicipes | crustacea |
| C.virginica | XP_022329852.1 | Crassostrea virginica | mollusca |
| M.galloprovincialis | VDI74315.1 | Mytilus galloprovincialis | mollusca |
| C.elegans | NP_500149.1 | Caenorhabditis elegans | nematoda |
| Auanema sp. | CAH6628979.1 | Auanema sp. JU1783 | nematoda |
| H.contortus | CDJ85118.1 | Haemonchus contortus | nematoda |
| A.tenebrosa | XP_031555938.1 | Actinia tenebrosa | cnidaria |
| A.millepora | XP_029211704.2 | Acropora millepora | cnidaria |
| D.gigantea | XP_028398530.1 | Dendronephthya gigantea | cnidaria |
| Xenia sp. | XP_046847875.1 | Xenia sp. Carnegie-2017 | cnidaria |
| H.vulgaris | XP_047136116.1 | Hydra vulgaris | cnidaria |
| Z.mays | NP_001169273.1 | Zea mays | plant |
| T.aestivum | XP_044456423.1 | Triticum aestivum | plant |
| B.napus | XP_013728150.2 | Brassica napus | plant |
| L.albus | KAE9606436.1 | Lupinus albus | plant |
| C.rubella | XP_006290125.2 | Capsella rubella | plant |
| L.lanzarotensis | XP_022629229.1 | Lachancea lanzarotensis | fungi |
| Z.parabailii | AQZ17865.1 | Zygosaccharomyces parabailii | fungi |
| T.globosa | XP_037141659.1 | Torulaspora globosa | fungi |
| K.africana | XP_003958391.1 | Kazachstania africana | fungi |
| C.glabrata | KAH7582295.1 | Candida glabrata | fungi |
| S.cerevisiae | AJQ13221.1 | Saccharomyces cerevisiae | yeast |
| D.bacterium | MBI1948579.1 | Deltaproteobacteria bacterium | bacteria |
| Burkholderia sp. | WP_226115822.1 | Burkholderia sp. AU33545 | bacteria |
| B.cosmicum | WP_143845348.1 | Bradyrhizobium cosmicum | bacteria |
| R.rhenobacensis | WP_184254289.1 | Rhodopseudomonas rhenobacensis | bacteria |
| Isosphaera sp. | MBA4064840.1 | Isosphaera sp. | bacteria |
| P.bacterium | MBA4190599.1 | Planctomycetaceae bacterium | bacteria |
| M.bacterium | MBK7077015.1 | Myxococcales bacterium | bacteria |
| G.bacterium | OGS99019.1 | Gallionellales bacterium | bacteria |
| R.bacterium | SEP49348.1 | Rhodospirillales bacterium | bacteria |
| C.eustigma | GAX84131.1 | Chlamydomonas eustigma | algae |
| C.sorokiniana | PRW59620.1 | Chlorella sorokiniana | algae |
| Pedinophyceae sp. | CAG9462817.1 | Pedinophyceae sp. YPF-701 | algae |
| Scenedesmus sp. | KAF8054979.1 | Scenedesmus sp. PABB004 | algae |
| Trebouxia sp. | KAA6424393.1 | Trebouxia sp. A1-2 | algae |
| M.conductrix | PSC74986.1 | Micractinium conductrix | algae |
| V.africanus | GIL60588.1 | Volvox africanus | algae |

